# Precision genome editing using synthesis-dependent repair of Cas9-induced DNA breaks

**DOI:** 10.1101/161109

**Authors:** Alexandre Paix, Andrew Folkmann, Daniel H Goldman, Heather Kulaga, Michael Grzelak, Dominique Rasoloson, Supriya Paidemarry, Rachel Green, Randall Reed, Geraldine Seydoux

## Abstract

The RNA-guided DNA endonuclease Cas9 has emerged as a powerful new tool for genome engineering. Cas9 creates targeted double-strand breaks (DSBs) in the genome. Knock-in of specific mutations (precision genome editing) requires homology-directed repair (HDR) of the DSB by synthetic donor DNAs containing the desired edits, but HDR has been reported to be variably efficient. Here, we report that linear DNAs (single and double-stranded) engage in a high-efficiency HDR mechanism that requires only ∼35 nucleotides of homology with the targeted locus to introduce edits ranging from 1 to 1000 nucleotides. We demonstrate the utility of linear donors by introducing fluorescent protein tags in human cells and mouse embryos using PCR fragments. We find that repair is local, polarity-sensitive, and prone to template switching, characteristics that are consistent with gene conversion by synthesis-dependent strand-annealing (SDSA). Our findings enable rational design of synthetic donor DNAs for efficient genome editing.

**Significance:** Genome editing, the introduction of precise changes in the genome, is revolutionizing our ability to decode the genome. Here we describe a simple method for genome editing that takes advantage of an efficient mechanism for DNA repair called synthesis-dependent strand annealing. We demonstrate that synthetic linear DNAs (ssODNs and PCR fragments) with ∼35bp homology arms function as efficient donors for SDSA repair of Cas9-induced double-strand breaks. Edits from 1 to 1000 base pairs can be introduced in the genome without cloning or selection.

## Introduction

Precision genome editing begins with the creation of a double-strand break (DSB) in the genome near the site of the desired DNA sequence change (“edit”) (1). Generation of targeted DSBs has been greatly accelerated in recent years by the discovery of CRISPR-Cas9, a programmable DNA endonuclease that can be targeted to a specific DNA sequence by a small “guide” RNA (crRNA) (2).

DSBs are lethal events that must be repaired by the cell’s DNA repair machinery. DSBs can be repaired via imprecise, non-homology-based repair mechanisms, such as non-homologous end-joining (NHEJ), or by precise, homology-dependent repair (HDR) (3). HDR utilizes DNAs that contain homology to sequences flanking the DSB (termed homology arms) to template the repair. If a synthetic “donor” DNA containing the desired edit is available when the DSB is generated, the cellular HDR machinery will use the donor DNA to repair the DSB and the edit will be incorporated at the targeted locus (1). Several studies have reported that single-stranded oligonucleotides (ssODNs) can be used to introduce short edits (<50 bases) ((4) and references therein). ssODNs that target the DNA strand that is first released by Cas9 after DSB generation have been reported to perform best (5). This strand preference, however, has only been tested for small edits near the DSB and has not been noticed at all loci (4). Edits at a distance from the DSB (>10 bp) are recovered at lower frequencies (4, 6). Recovery of large edits (such as GFP knock-ins) has also been reported to be inefficient, requiring large plasmid donors with long (>500nt) homology arms or selection markers to recover the rare edits (3). Large insertions have been obtained through non-homologous or micro-homology-mediated end joining reactions (NHEJ and MMEJ), but these approaches require simultaneous Cas9-induced cleavage of donor and target DNAs (7-13).

We documented previously that, in *C. elegans*, HDR can be very efficient provided that the donor DNAs are linear (14). Linear donors do not appear to integrate at the DSB, but instead are used as templates for DNA synthesis, as in the synthesis-dependent strand annealing (SDSA) model for gene conversion (Figure 1) (15). In SDSA, the DSB is first resected to yield 3’ overhangs on both sides of the DSB (Figure 1A). The 3’ overhangs invade the donor and are extended by DNA synthesis copying donor sequences (Figure 1B). Bridging of the DSB is completed when the newly synthesized strands withdraw from the donor and anneal back to each other at the locus (Figure 1C). In *C. elegans*, donors for SDSA can be single (ssODNs) or double-stranded (PCR fragments), and require only short homology arms (∼35 bases) to engage the DSB. The repair process is sensitive to insert size and prone to template switching, where synthesis can “jump between two overlapping donors (14). In human cells, SDSA has been proposed as a repair mechanism for ssODNs (4, 16), but not for double-stranded donors, which are thought to participate in a different HDR pathway (16, 17). Here, we investigate the sequence requirements for linear donors to engage the DSB repair machinery in human cells. Our findings suggest that ssODNs and PCR fragments both engage in a SDSA-like type of gene conversion, and we demonstrate the utility of PCR fragments to create fluorescent protein knock-ins in human cells and mouse embryos. Our findings suggest simple donor DNA design principles to maximize editing efficiency.

**Figure 1:**
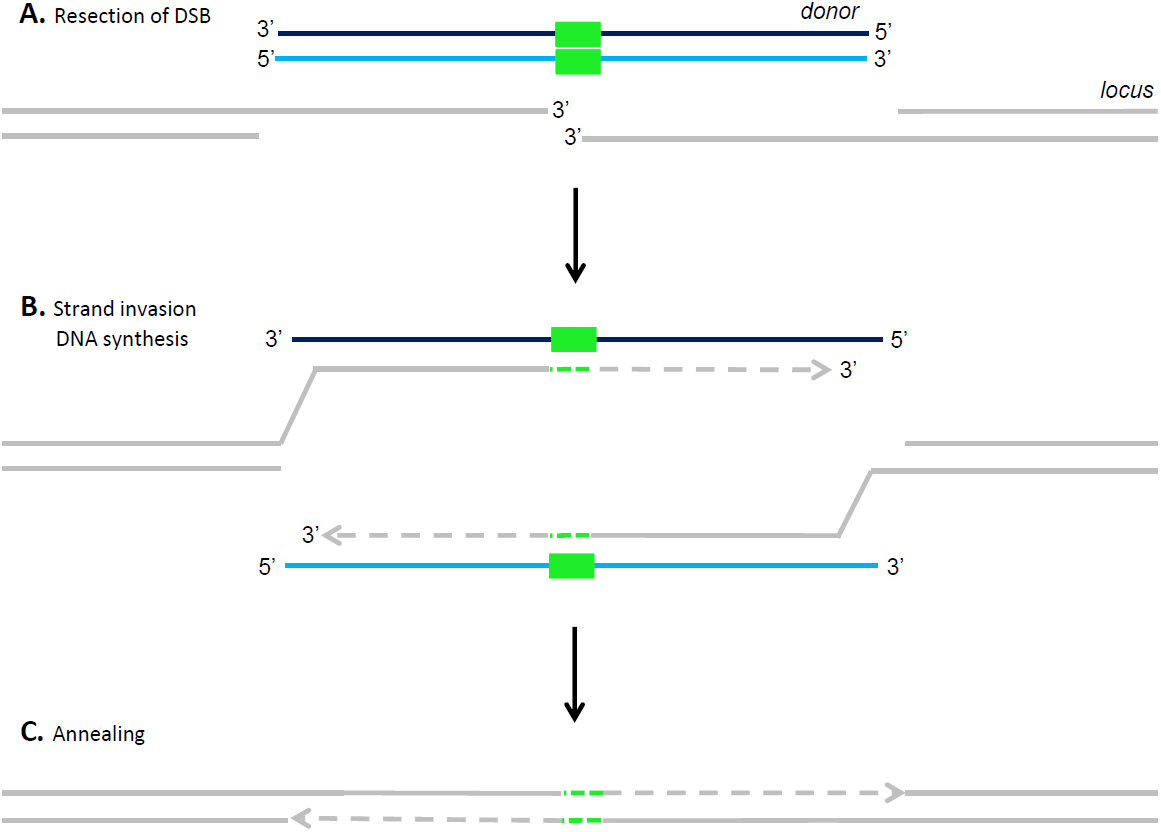
Synthesis-dependent strand annealing (SDSA) model for gene conversion. Diagrams showing SDSA model for gene conversion (after (15)). Each line corresponds to a DNA strand: locus - grey, donor - blue, edit - green. Dotted lines represented newly synthesized DNA. **A.** DSB is resected creating two 3’overhangs on each side of the DSB. **B.** Strand invasion / DNA synthesis: The overhangs pair with complementary strands in the donor and are extended by DNA synthesis. **C.** Annealing: The newly synthesized strands withdraw from the donor and anneal back at the locus. Ligation seals the break.

## Materials and Methods

### Detailed results, sequences and solutions

Table S1 lists all experiments, including detailed conditions and results of experimental replicates. Table S2 to S5 lists sequences of linear donors, plasmids, PCR primers and cr/sgRNAs, respectively. Position of the cr/sgRNAs on the loci targeted in this study can be found in Figure S1.

### Repair templates, Cas9, cr/tracrRNAs and plasmids for cell culture

ssODNs (ultramers) and PCR primers where ordered from IDT and reconstituted at 50uM and 100uM respectively in water. For the Illumina sequencing experiment shown in Figure 7B, ssODNs and primers were ordered PAGE purified. PCR fragment donors were synthesized as described in (18).

Cas9 protein was purified as described in (19). crRNAs and tracrRNA were ordered from IDT and reconstituted in 5mM Tris-HCl pH7.5 at 130uM. *PYM1* sgRNA cloning was cloned as described in (20). Plasmids containing repair templates were made using gBlock gene fragments (IDT) and InFusion cloning kit (Clontech), and purified using Qiagen mini-prep kit and eluted in H2O.

### Cas9 RNP nucleofection

With the exception of experiments at the *PYM1* locus (see below), all experiments in this study used Cas9 RNP delivery (21). Nucleofections using Cas9 RNP were performed as described (22). HEK293T cells or HEK293T cells expressing a truncated GFP (GFP1-10) (23) were grown to 50-75% confluency, trypsinized, pelleted and resuspended at 800000 cells / 80ul of PBS. Just before nucleofection, PBS was replaced with 80ul of Nucleofection kit V (Lonza). 40ul of Cas9 RNP mix (see below) was added to the cells in suspension in Nucleofector kit V and processed using an Amaxa Nucleofector 2b machine (Lonza) using the A023 program. Cells were transferred to culture media and analyzed for fluorescence 3 days after.

The Cas9 RNP mix contains: 6.5uM of crRNA and tracrRNA, 1.6ug/ul of Cas9, a variable concentration of repair templates (see Table S1 for details), 10.4% Glycerol, 131mM KCl, 5.2mM Hepes, 1mM MgCl2, 0.5mM Tris-HCl, pH7.5.For sequencing of GFP edits at the *Lamin A/C* locus, cells were sorted (at the JHU Ross Flow Cytometry Core Facility) for GFP signal and cloned in 96 wells plates for genotyping or pooled in a 6-well plate for microscopy analysis. Single cell clones were lysed using QuickExtract DNA Extraction Solution (Epicentre) and genotyped by PCR using Phusion taq (NEB) with genomic primers outside of the HDR fragment. PCR products were analyzed on agarose gel and sequenced (see Figures S6 and S7).

### Cas9 plasmid transfections

For experiments at the *PYM1* locus, Cas9 and the sgRNA were delivered on plasmids. HEK293T cells were grown to 50-75% confluency in 6 wells plate (with 2ml of culture media per wells). 10.8ul of Cas9 plasmid mix (containing 3.6ul of X-tremeGENE 9 DNA Transfection Reagent from Roche, 892ng of plasmid pX458 containing *PYM1* sgRNA and 3.24pmol of repair template) was added to 120ul of optiMEM glutaMAX media (ThermoFisher), incubated for 15min at room temperature, and next added to the cells. 48h after transfection, cells were sorted for GFP signal (to select for cells that received pX458) and grown out as single cell clones. The single cell clones were lysed and genotyped by PCR. PCR products were directly analyzed on agarose gel or mix with *EcoR1* (NEB) and the corresponding Restriction Enzyme (RE) buffer, digested over-night and analyzed on agarose gel.

### Cytometer analysis

For each experiment, 5,000 to 10,000 cells were analyzed using a Guava EasyCyte 6/2L (Millipore) cytometer. Cells were scored as GFP+ if they exhibited a higher signal than 99.5% of non-transfected control cells.

HEK293T (GFP1-10) cells exhibit a higher basal green fluorescence than wild-type HEK293T cells. Cytometer analysis could not be performed on these cells for GFP11-tagged Lamin A/C and SMC3. For those experiments, as well as for RFP tagging, cells were analyzed by fluorescence microscopy and scored manually (see below).

### Microscopy

Cells were fixed in 4% PFA and mounted with DAPI. Cells were imaged using a confocal microscope with a 63X objective. > 50 fields of cells (>1000 cells) were selected in the DAPI channel, photographed, and analyzed for GFP or RFP expression manually.

### PCR amplicons for Illumina sequencing

HEK293T (GFP1-10) were nucleofected with different combinations of repair ssODNs (Fig 7B, Table S1). To control for possible template-switching during PCR amplification, we also introduced single donors (wild-type or mutant) in two separate cell populations and combined the cells during PCR amplification. 60h after nucleofection, cells were trypsinized, washed in PBS, and 500000 cells were lysed in 40ul of QuickExtract DNA Extraction Solution. 40ul of H2O was added to each lysis. A total of 6ul of DNA from each experiments were PCR amplified using Phusion Taq and the primer 390 (Forward, in the left end of the insert) and the primer 1849 (Reverse, in the *Lamin A/C* locus downstream of the right HA of the ssODN used for repair) for 10 cycles at 68.5C (see Table S4 for primer sequences). After 10 PCR cycles, no band could be detected on agarose gel and ethidium bromide staining. Each PCR reaction was purified using Qiagen Minelute columns and eluted in 10ul of H2O. 2ul of each PCR were amplified using Phusion taq at 65C for 20 cycles. PCR reactions did not reach an amplification plateau with this number of cycles. The PCR reactions were performed using primers 1928 (Forward, containing the Illumina sequence and annealing in the same region than primer 390) and Reverse primers containing the Illumina sequence and a specific barcode. The Illumina reverse primers anneal with the *Lamin A/C* locus just upstream of primer 1849 and downstream the right HA of the ssODN used for repair.

PCR amplicons were purified on a 10% non-denaturing TBE/PAGE gel and the band corresponding to the PCR product was cut from the gel, eluted over-night, and precipitated with isopropanol. After resuspension, sample concentrations were quantified on a bioanalyzer, and the barcoded samples were pooled to a concentration of 0.4uM per sample in 10 ul. This sample was submitted to the Johns Hopkins School of Medicine Genetics Resources Core Facility for 250 cycle paired-end sequencing on an Illumina MiSeq instrument.

### Illumina Sequencing analysis

After de-multiplexing of barcoded samples, the 3’ adaptor and all downstream nucleotides were trimmed from the forward reads using Cutadapt (<u>http://journal.embnet.org/index.php/embnetjournal/article/view/200</u>), and the resulting sequences were mapped to the insert + *Lamin A/C* locus using Bowtie 2 (24). After removing reads that did not fully map to the template and low-quality reads (Q score less than 35; error probability of 0.00032), sequences were parsed for template switching. To score template switches, we evaluated sequencing reads at diagnostic positions and determined whether each position matched the sequence of the wild-type or mutated template. Reads with a diagnostic nucleotide that did not match either the wild-type or mutated template were discarded. Because the PCR control sample contained a mixture of the fully wild-type and fully mutated templates, we used the first diagnostic position (from the right side of the insert) only as an “anchor” to determine the initial identity of the template; this position was not used to score switching. Thereafter, whenever two or more contiguous diagnostic nucleotides indicated a switch in template identity, we scored this as a switch. For the control sample in which both templates were wild-type, we used the “1/6” mutated template for comparison, to determine the rate of false-positive switches in the assay. Because the PCR control experiment was performed with the wild-type and “1/6” mutated template (Table S1), we also used the “1/6” mutated template for scoring switches in this sample. See Table S6 for details.

### Cas9 RNP injection in mouse zygotes

All mouse experiments were carried out under protocols approved by the JHU animal care and use committee.

The PCR fragment donor was synthesized as described in (18). The plasmid donor was generated using a gBlock and restriction enzyme cloning, and purified by Qiagen midi-prep kit and eluted in injection buffer (10 mM Tris-HCl, pH 7.5, 0.1 mM EDTA). Pronuclear injections of zygotes (from B6SJLF1/J parents (Jackson labs)) was performed by the JHU Transgenic facility at a final concentration: 30ng/ul Cas9 protein (PNABio), 0.6uM each of crRNA/TracrRNA (Dharmacon) and PCR donor (3ng/ul or 5ng/ul) or plasmid donor (10ng/ul). The Cas9 protein, crRNA, tracrRNA were combined from stocks at 1000ng/ul, 20uM, 20uM respectively and incubated at 4C for 10 minutes. Then injection buffer was added to dilute to the final working concentrations above (Table S1) along with repair vector or fragment. The solution was microcentrifuged 5 min at 13000xg and the solution used for injection. Pups were genotyped using genomic primers immediately outside of the PCR donor sequence, or using one primer in mCherry and one upstream of the 483 bp homology arms in the case of the plasmid donor.Genomic DNA from all pups was also subjected to PCR amplification with internal mCherry specific primers to identify random insertions of the donor template (locus-specific mCherry negative/internal mCherry product positive). We identified 7 pups (11%, out of 60 pups without mCherry insertion at the *Adcy3* locus) with potential transgenic insertions of the PCR fragment at other undetermined loci. In contrast, we identified no transgenics (0%, out of 20 pups without mCherry insertion at the *Adcy3* locus) when using the plasmid donor.

## Results

### PCR fragments with short homology arms are efficient donors for genome editing in HEK293T cells

In human cells, ssODNs and plasmids are most commonly used as donors for genome editing (3). To determine whether PCR fragments can also function as donors, we attempted to knock-in GFP at three loci using PCR fragments amplified from a GFP-containing plasmid. The GFP-coding insert (714 bp) was amplified using hybrid primers containing sequences to target GFP and ∼35 bp locus-specific homology arms (HAs). The HAs were designed to insert GFP in frame with the target open-reading frame 0, 11 and 5 bp away from a Cas9 cleavage site in the *Lamin A/C, RAB11A*, and *SMC3* ORFs, respectively (Figures 2 and S2). The PCR fragments (0.33-0.21uM) and *in vitro*-assembled Cas9-guide RNA complexes were introduced by nucleofection into HEK293T cells without selection as in (22). The efficiency of GFP integration was examined 3 days later by cytometer or fluorescence microscopy (see Material and Methods). We obtained 14.9%, 17.5% and 13.9% GFP+ positive cells for the *Lamin A/C, RAB11A* and *SMC3* loci, respectively (Figure 2B and S2B). In each case, the cells expressed GFP in a pattern consistent for the targeted ORF (Figures 2D and S2C).

**Figure 2:**
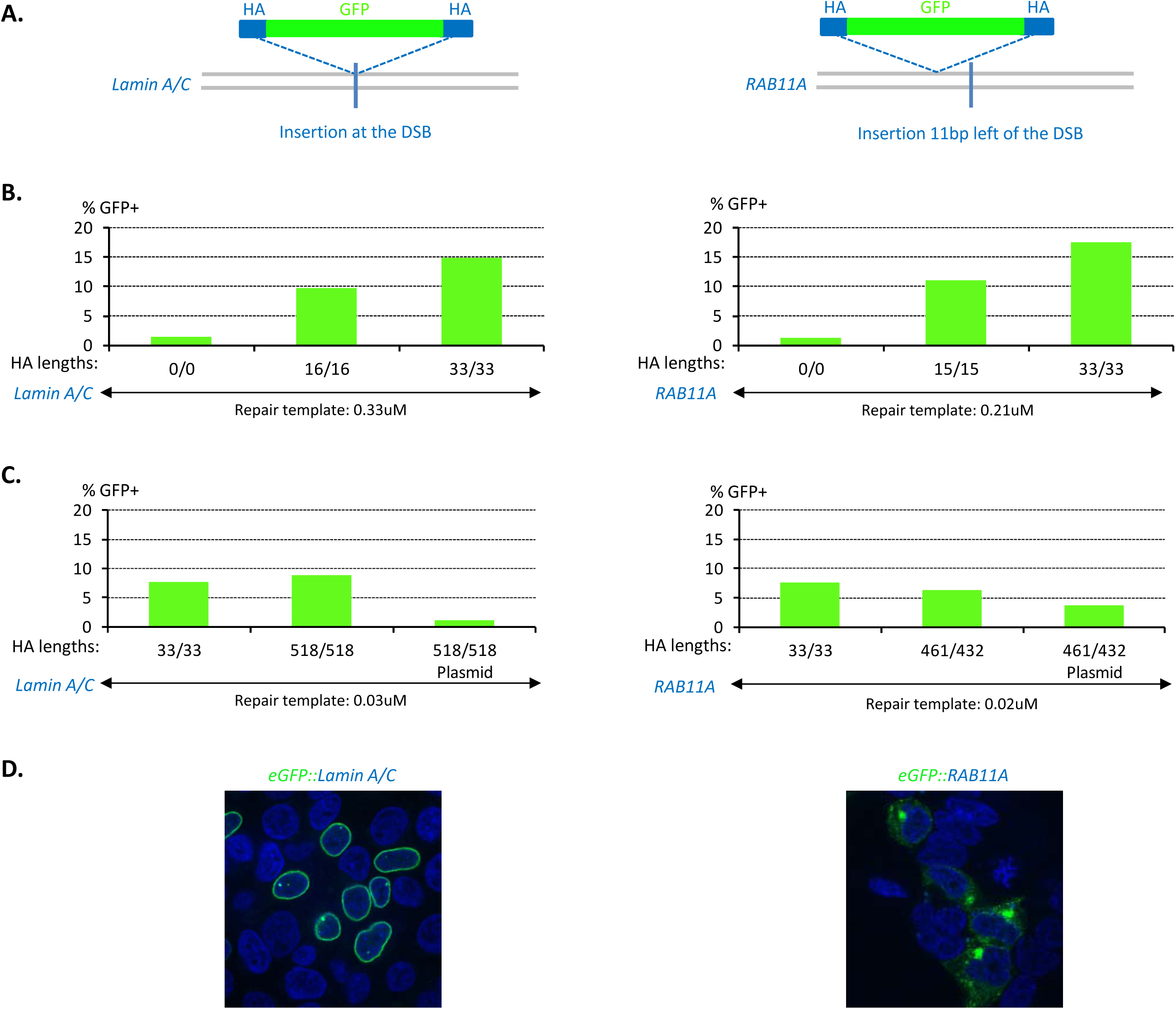
PCR fragments with short homology arms are efficient donors for GFP insertion. **A.** Diagrams showing PCR donors for GFP insertion at the *Lamin A/C* and *RAB11A* loci. Locus - grey, GFP - green, HA (Homology Arm) - blue, DSB - vertical line. GFP was inserted at the DSB in *Lamin A/C* and 11 bp upstream of the DSB in *RAB11A* locus. **B.** Graphs showing % of GFP+ cells obtained with PCR donors with HAs of the indicated lengths (33/33 refers to a right HA and a left HA, each 33 bp long). Insert size in all cases was 714 bp. PCR fragments were nucleofected in HEK293T cells at the concentration indicated and cells were counted by flow cytometer 3 days later. For this and all other figures, see Table S1 for details. **C.** Graphs showing % of GFP+ cells obtained with PCR or plasmid donors with HAs of the indicated lengths. Insert size in all cases was 714 bp. PCR fragments were nucleofected in HEK293T cells at the concentration indicated and cells were counted by flow cytometer 3 days later. **D.** Confocal images of cells 3 days after nucleofection. GFP: green, DNA: blue. The GFP subcellular localizations are as expected for in frame translational fusions.

Reducing the molarity of the PCR fragments by 10-fold reduced efficiency by ∼1/2 (Compare Figures 2B and 2C). Increasing the length of the homology arms to 500 bp did not increase editing efficiency significantly, even when controlling for the reduced molarity of the longer PCR fragments (Figure 2C). Reducing the length of the homology arms to ∼15 bp, however, decreased efficiency (Figure 2B). PCR fragments with no homology arms or homology arms for a locus not targeted by Cas9 yielded GFP+ edits near background levels (Figures 2, S2, S3 and Table S1). Plasmid donors with ∼500 bp homology arms also performed poorly (Figure 2C) as reported previously (7). We conclude that PCR fragments function as efficient donors in HEK293T cells, performing similarly to ssODNs, and better than plasmids with much longer homology arms. Because ∼35 bp homology arms are convenient to introduce by PCR amplification, we used that length for subsequent experiments. 30-40 nt homology arms have also been reported to be optimal for ssODNs (4).

### Editing efficiency is sensitive to insert size

To test the effect of insert size on editing efficiency, we added varied sizes of DNA sequence to the GFP insert. For ease of synthesis and to maintain equimolar amounts of donor DNAs, we introduced donor fragments at the same low molarity (0.12uM). We found that inserts beyond 1kb performed very poorly, yielding less than 0.5% edits (Figure 3A). By varying the size of the homology arms, we found that the size of the insert, and not the overall size of the donor DNA, determines editing efficiency. An 1188 bp donor (714 bp insert with two 237 bp HAs) performed as well as a 780 bp donor with the same size insert and 33 bp HAs (10% versus 11% edits, Figure 3A). The 1188 bp donor, however, performed much better than a 1188 bp donor with a longer insert (1122 bp) and 33 bp HAs (10% versus 0.5% edits, Figure 3A).

**Figure 3:**
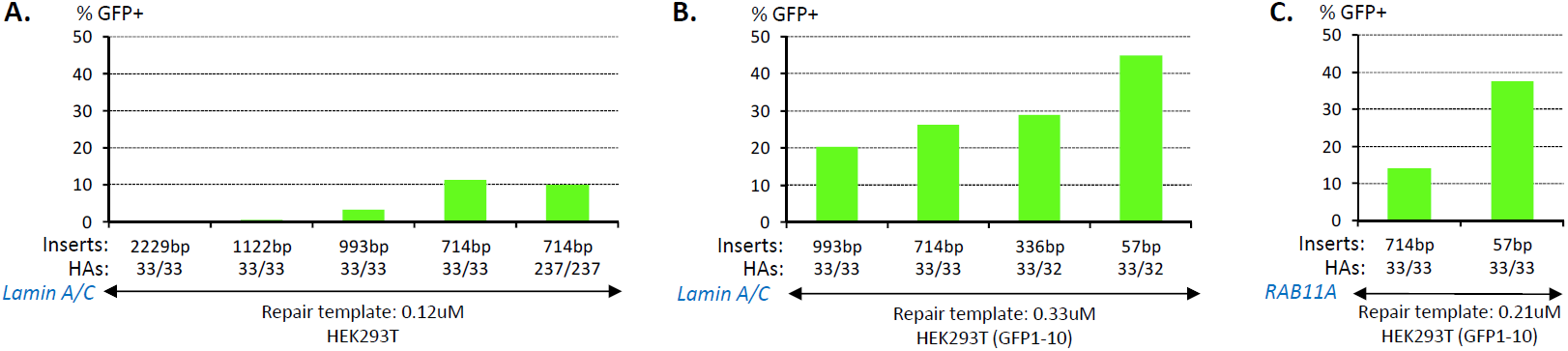
Editing efficiency increases with decreasing insert size. A.Knock-in of donors containing full-length GFP at the *Lamin A/C* locus. PCR fragments were nucleofected in HEK293T cells at the concentration indicated and cells were counted by microscopy 3 days later. HAs were 33/33, except for the last column where HAs were 237/237. **B.** Knock-in of donors containing full-length GFP or GFP11 at the *Lamin A/C* locus. PCR fragments were nucleofected at the concentration indicated in HEK293T (expressing GFP1-10) and cells were counted by microscopy 3 days later. HAs were 33/32 (57 and 336 bp inserts) or 33/33 (714 and 993 bp inserts). **C.** Knock-in of donors containing full-length GFP or GFP11 at the *RAB11A* locus (11 bp upstream of DSB). PCR fragments were nucleofected at the concentration indicated in HEK293T (expressing GFP1-10) and cells were counted by flow cytometer 3 days letter. HAs were 33/33.

To test whether decreasing insert size below the size of GFP would increase editing efficiency, we took advantage of the split-GFP system (22, 23). In this system, the 11^th^ beta-strand of GFP (57 bp, GFP11) is knocked-in in cells expressing a complementary GFP fragment (GFP1-10). We generated PCR products containing the GFP11 insert and ∼35 bp HAs and introduced these at 0.33uM. We obtain 44.9% edits at the *Lamin A/C* locus (Figure 3B) and 37.4% at the *RAB11A* locus (Figure 3C). A donor with no homology arms yielded only 1.3% edits (Figure S3B). Again, we found that increasing insert size reduced efficiency, down to 20% for a 993 bp insert (Figure 3B). We conclude that dsDNAs engage in an efficient repair process that requires only 35 bp homology arms, but favors relatively short inserts (<1kb at the molarities tested here).

### Repair is a polarity-sensitive process

In the SDSA repair model, when using a double-stranded donor, strand invasion can be initiated on either side of the DSB, since both strands in the donor are available for pairing (Figure 1) (15). In the case of single-stranded donors (ssODNs), however, the model predicts that repair will initiate only on the right or left side of the DSB depending on the polarity of the ssODN. To test this prediction, we designed ssODNs with a GFP11 insert and only one HA targeting either the left or right side of the Cas9-induced DSB in *Lamin A/C* and *RAB11A* (Figure 4). ssODNs with only one homology arm were able to function as donors, presumably because NHEJ can repair the gap on the side with no HA (see discussion). As predicted by the model, the polarity of the ssODN had a profound effect on editing efficiency. Editing efficiency was highest with ssODNs that could anneal to a complementary 3’ end at the DSB (Figure 4). These observations are consistent with a replicative repair process initiated by pairing between 3’ overhangs at the DSB and the HAs on the donor.

**Figure 4:**
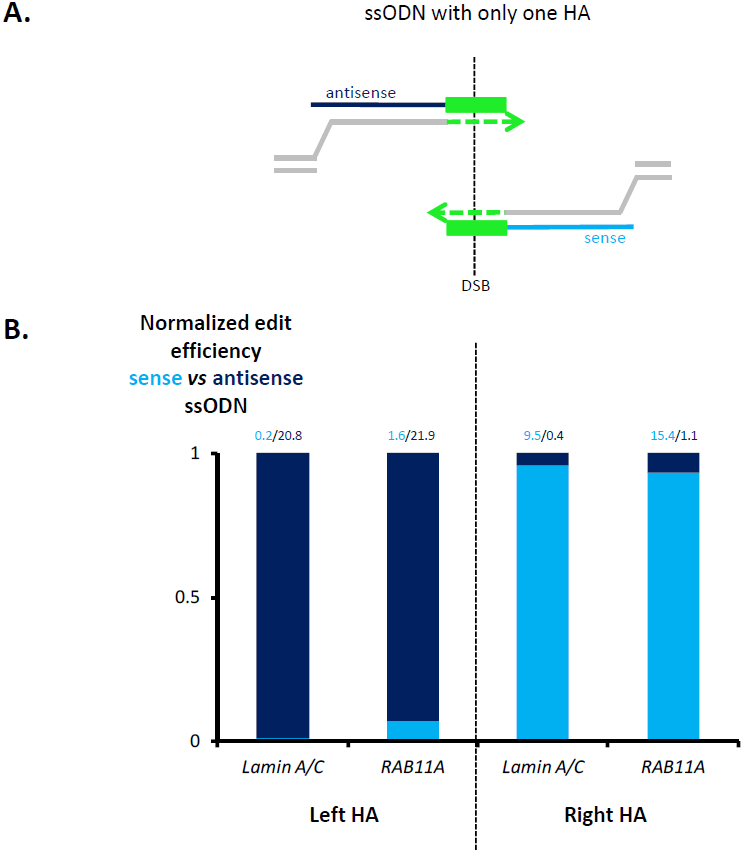
Repair is a polarity-sensitive process. **A.** Diagrams showing pairing of ssODNs (antisense: dark blue, and sense: light blue) with resected ends of the locus (grey). Dotted lines denote DNA synthesis as in Figure 1. The ssODNs have only one 32-33 bp HA and a 126 bp insert (containing a combination of 3xFlag and GFP11 – green). **B.** Normalized efficiency of sense *vs* antisense ssODNs containing only one HA corresponding to sequence on left or right side relative to DSB. The polarity that allows pairing between the ssODN and resected ends (as shown in diagram in A) is favored. Numbers on top of each column indicate the % of GFP+ cells determined by microscopy (*Lamin A/C*) or flow cytometer (*RAB11A*).

### Recoding of sequences between the DSB and the edit increases recovery of distal edits

Editing efficiency is known to decrease with increasing distance between the edit and the DSB (6). This observation is also consistent with replicative repair, which predicts that synthesis that generates sequence complementary to the other side of the DSB will promote “premature annealing” back to the locus before synthesis of the edit (Figure 5). To test this prediction directly, we designed an ssODN donor with two inserts: a proximal insert (restriction enzyme site) at the DSB in the *PYM1* locus and a distal insert (3xFlag) 23 bases away from the DSB. Each insert was flanked by an HA targeting the *PYM1* locus (Figure 5A). We generated 63 single cell clones and genotyped the *PYM1* locus by PCR and Sanger sequencing (see Material and Methods). 46% of the clones contained only the proximal edit and 12.6% contained both the proximal and distal edits. The finding that ∼80% of the edits contained only the proximal edit is consistent with “premature” annealing using sequence between the two edits. To test this hypothesis, we mutated 7 bases in the 23 bases region separating the proximal and distal edit. The mutations were designed to reduce homology with the locus while preserving coding potential (Figure 5A). This partial recoding reduced the frequency of proximal edit-only clones to 10.3% and increased the frequency of proximal+distal edits to 25.8%. We conclude that sequences on the donor that span the DSB can trigger premature annealing and prevent the incorporation of distal edits.

**Figure 5:**
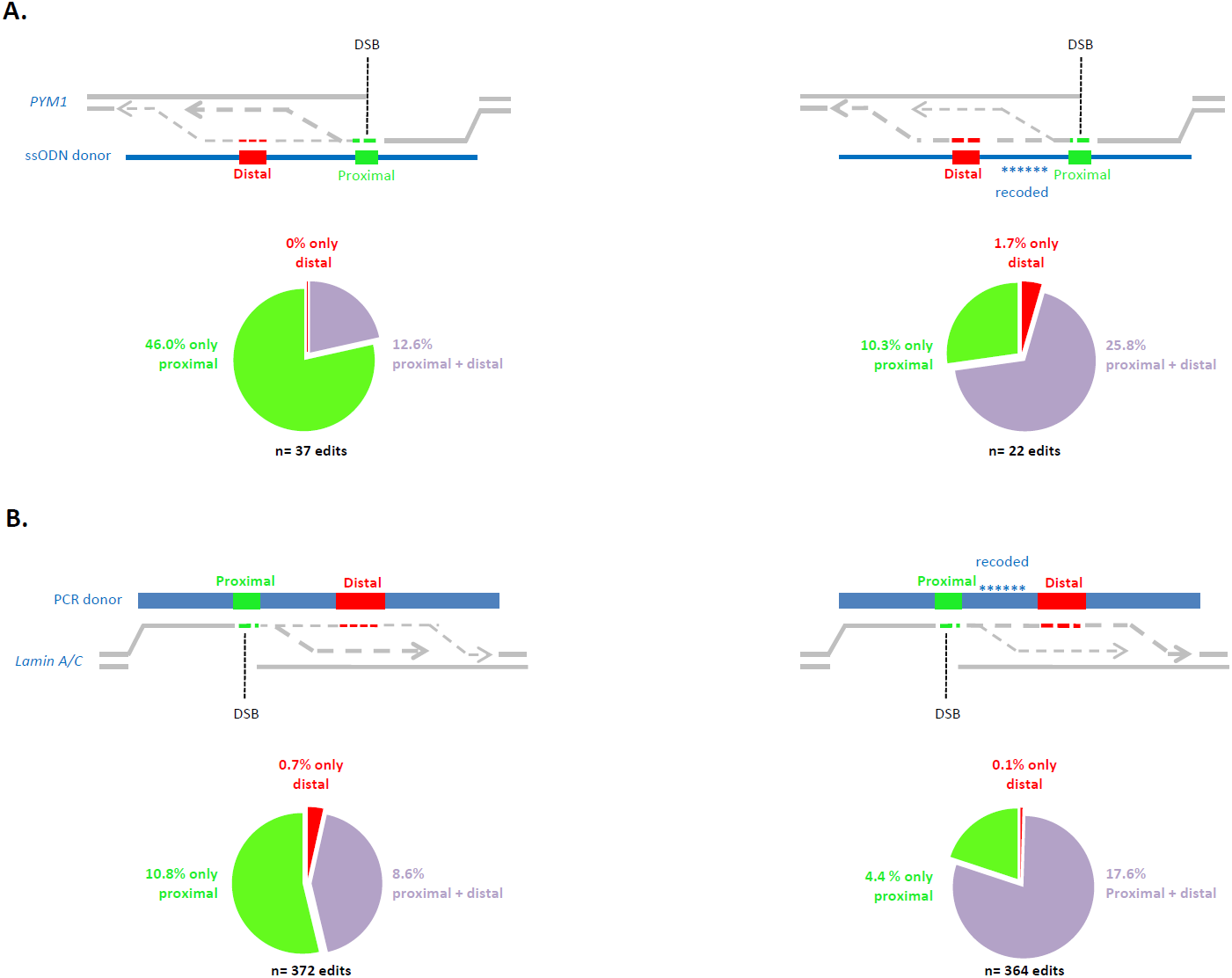
Recoding of sequences between the DSB and the edit increases recovery of distal edits. **A.** Schematics showing possible interactions between resected locus (grey, *PYM1* locus) and ssODNs coding for proximal and distal edits. Dotted lines denote DNA synthesis. Proximal edit (green, Restriction Enzyme RE site) is 1 bp to the right of the DSB and distal edit (red, 3xFlag) is 23 bp to the left of the DSB. Asterisks denote silent mutations in the region between the distal and proximal edits. Circles show the relative frequency of cells containing both edits (purple), only the proximal edit (green), or only the distal edit (red). % refers to the percent of each edit types among all the cell clones analyzed analyzed by PCR genotyping (size shift) and RE digestion. N refers to the total number of edited cell clones. **B.** Schematics showing possible interactions between resected locus (grey, *Lamin A/C* locus) and PCR fragments coding for proximal and distal edits. Dotted lines denote DNA synthesis. Proximal edit (green, GFP11) is at the DSB and distal edit (red, tagRFP) is 33 bp to the right of the DSB. Asterisks denote silent mutations in the region between the distal and proximal edits. Circles show the relative frequency of cells containing both edits (purple), only the proximal edit (green), or only the distal edit (red). Edits were determined by microscopy. % refers to the percent of each edit type among all cells analyzed by microscopy. N refers to the total number of edited cells.

To test whether internal homologies can also participate in the repair process in double-stranded donors, we performed a similar experiment with a PCR fragment designed to incorporate GFP11 at the DSB, and tagRFP 33 bases from the DSB in the *Lamin A/C* locus (Figure 5B). We recovered 10.8% GFP-only edits and 8.6% GFP-RFP double positives. Partial recoding of the sequence between GFP11 and tagRFP (by introducing 10 silent mutations) reduced the percent of GFP-only edits to 4.4% and raised the percent of GFP-RFP double positives to 17.6%. We conclude that internal homologies on double-stranded templates can also interact with the targeted locus. Since both polarities are present in double-stranded templates, internal sequences could participate in principle in both the initial invasion step and the annealing step back to the locus.

### Polarity of single-stranded donors affects incorporation of distal edits

Donors that contain edits designed to be inserted at a distance from the DSB have one HA that matches sequences immediately next to the DSB (proximal HA) and one HA at a distance from the DSB (recessed arm) (Figure 6A). We tested whether proximal and recessed arms initiate repair with similar efficiencies using a series of 22 ssODNs with inserts ranging from 0 to 41 nucleotides from the DSB at four loci (Figure 6). In all ssODNs, the sequence between the DSB and edit was partially recoded to increase the frequency of edit incorporation as described in the previous section. Strikingly, we observed an increasing bias for a particular polarity with increasing edit-to-DSB distance (Figure 6B). The favored polarity changed whether the edit (and recessed arm) was positioned to the left or right of the DSB (sense polarity when the edit is on the left side of the DSB, and antisense when the edit is on the right side). ssODNs with inserts close to the DSB did not show much polarity bias (Figure 6B). These findings are consistent with a replicative model for repair where proximal arms directly abutting the DSB are favored to initiate a round of DNA synthesis that will copy the edit. Recessed arms can be used for annealing back to the locus, although that process also appears to favor proximal arms. Even when using ssODNs with the correct polarity and recoding between the DSB and the edit, we still observed an inverse correlation between editing efficiency and edit-to-DSB distance (Figure S4). Finally, we note, that, unlike ssODN polarity, the polarity of the guide RNA used to create the DSB had only a minor effect, if any, on efficiency (Figure 6B).

**Figure 6:**
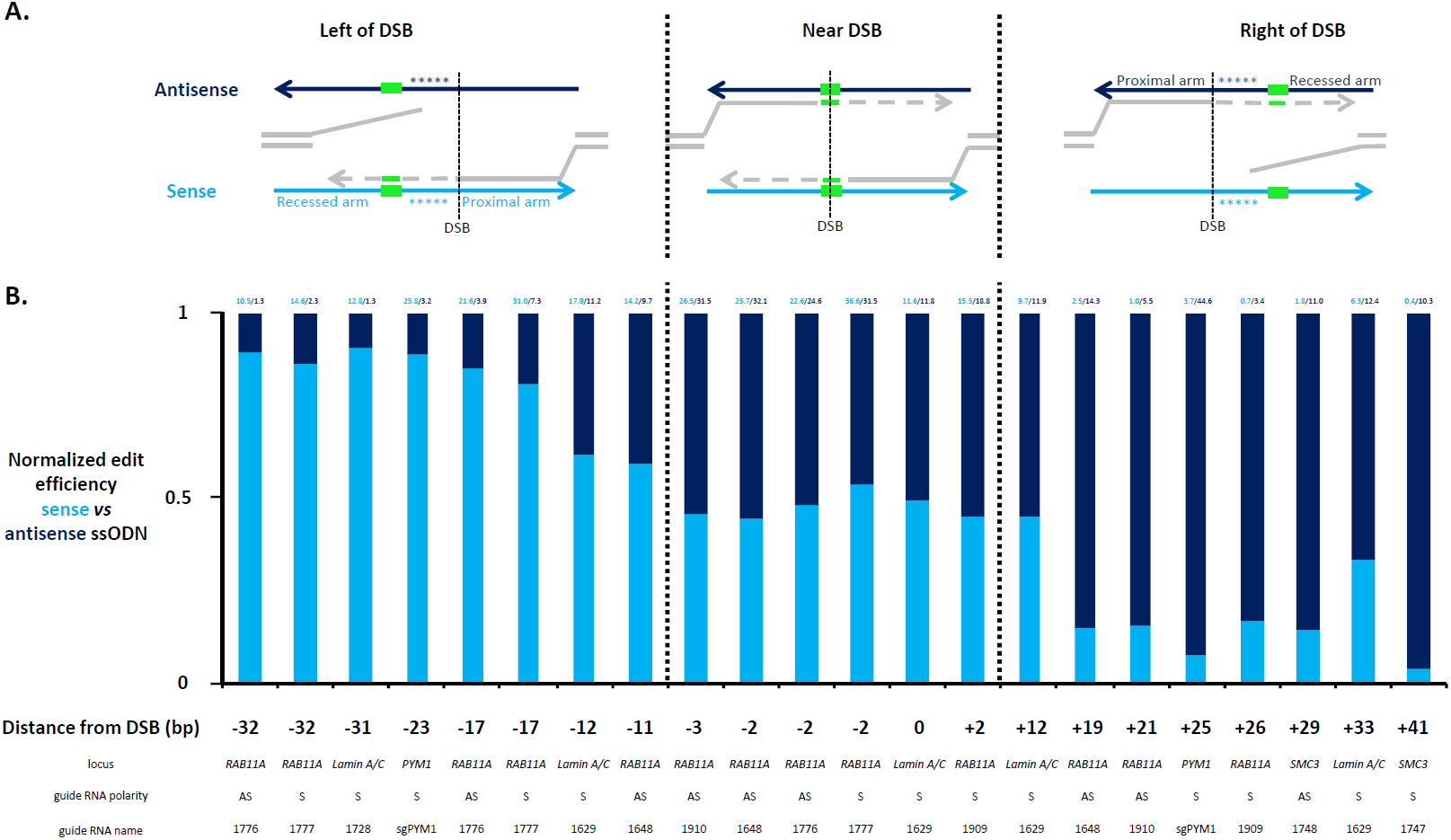
Polarity of ssODNs affects incorporation of distal edits. **A.** Schematics showing possible interactions between resected locus (grey) and ssODNs (light or dark blue depending on polarity) coding for a distal edit (green). Dotted lines denote DNA synthesis. Asterisks denote silent mutations in the region between the DSB and the distal edit. **B.** Normalized efficiency of sense *vs* antisense ssODNs with edits inserted to the left of the DSB, near the DSB, or to the right of the DSB. Distance from the DSB is indicated under each experiment. The locus and guide RNA and polarity are also indicated below. ssODN polarity has little effect on editing efficiency for proximal edits. In contrast, ssODN polarity has a large effect for distal edits. The favored polarity changes depending on whether the distal edit is positioned to the left or right of the DSB. Note that the favored ssODN polarity does not correlate with crRNA polarity (for example, first two columns in the graph show crRNAs 1776 and 1777 which cut at the same position but have opposite polarity). Experiments involving the *PYM1* locus were done on HEK293T that were cloned out and genotyped by PCR genotyping (size shift) for 3xFlag insertion (see Figure 5). All other experiments were performed on HEK233T (GFP1-10) cells that were directly scored for GFP+ by flow cytometer or microscopy 3 days after nucleofection. Numbers on top of each column indicate the overall % of edits. Note that overall frequency decreases with increasing distance from the DSB (see Figure S4).

**Figure 7:**
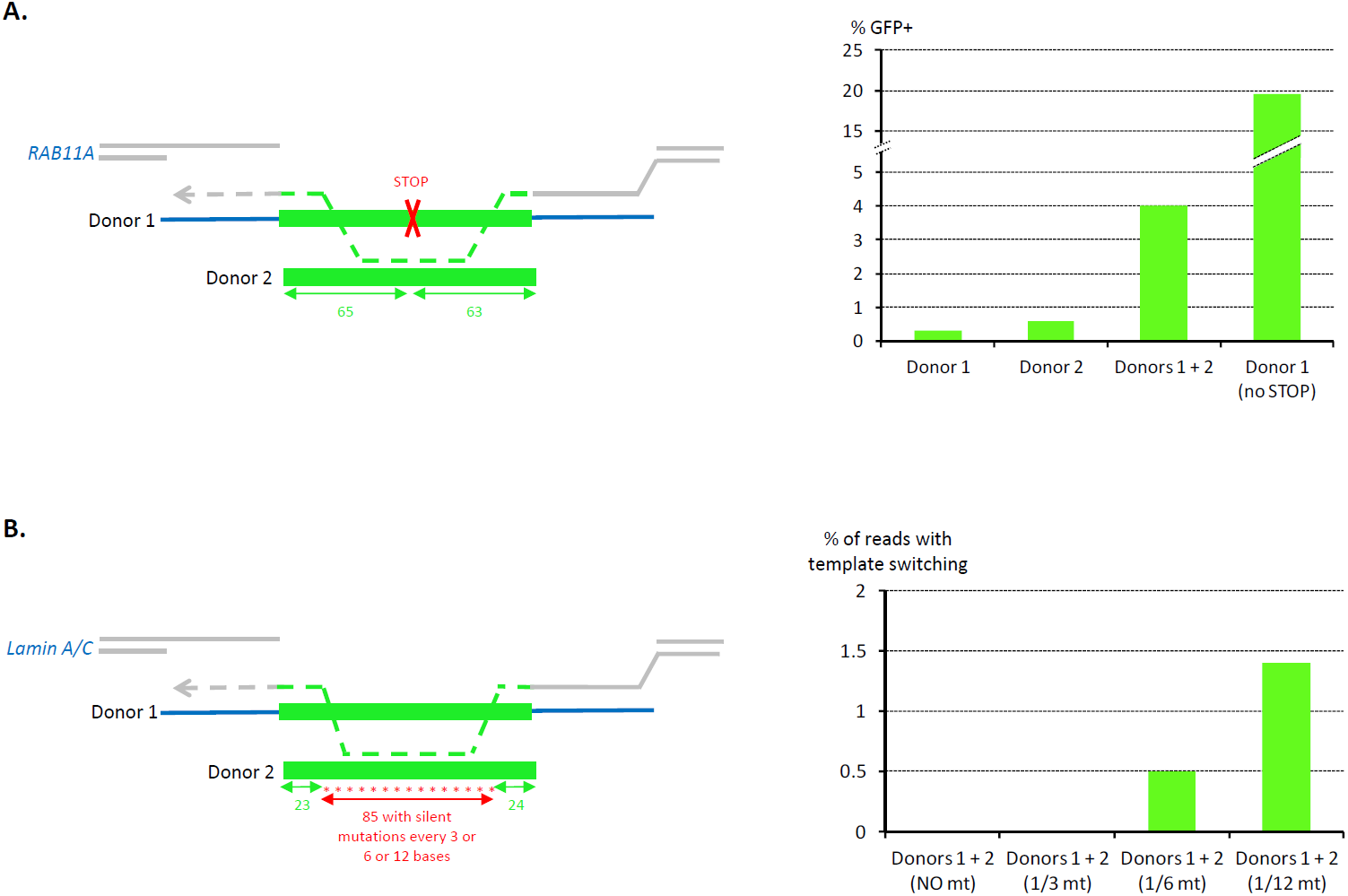
Repair is prone to template switching between donors. **A.** Schematics showing repair of a DSB at the *RAB11A* locus with two donors. Donor 1 contains GFP11 with a STOP codon and two HAs. Donor 2 contains GFP11 with no stop codon and no HA. Double arrows indicate identical sequence shared between the donors. Graphs showing the percent of GFP+ cells (Y axis, as determined by flow cytometer) for each donor combination (X axis). Each donor used separately gives % of GFP+ close to background whereas donor 1 + donor 2 gives GFP positives cells (4%, GFP expression confirmed by microscopy). As a reference, an ssODN identical to donor 1 but without the STOP codon gives 19.6% edits (discontinuous right column). B.Schematics showing repair of a DSB at the *Lamin A/C* locus with two donors. Donor 1 contains GFP11 and two HAs. Donor 2 contains a recoded GFP11 (stars) with no HA. Double arrows indicate identical sequence shared between the donors. In this experiment, the edits were amplified en masse by PCR using a locus-specific primer and an insert-specific primer and sequenced by Illumina sequencing (see Material and Methods). Graph showing the % of reads with evidence of template switching (Y axis) for each donor combination (X axis). Donor 1 + donor 2 without mutations and donor 1 + donor 2 with 1 mutations every 3 nucleotides (1/3) show no evidence of template switching (0%), whereas donor 1 + donor 2 (1/6) and donor 1 + donor 2 (1/12) show evidence of template switching (0.5% and 1.4% respectively). See Figure S5 and Table S6 for details.

### Repair is prone to template switching between donors

In *C. elegans*, we observed that sequential rounds of invasion and synthesis (“template switching”) can create edits that combine sequences from overlapping donors (14). To test whether template switching also occurs in human cells, we combined two donors in a single editing experiment. One donor was an ssODN with two HAs and an insert containing GFP11 with a STOP codon that prevents translation of the full-length fusion (Figure 7A). The second donor was an ssODN of the same polarity but with no HAs and no STOP codon (wild-type GFP11 insert). Consistent with template switching, we obtained 4% GFP+ edits when using both donors, compared to 0.3% and 0.6% GFP+ edits when using only the first or second ssODN, respectively (Figure 7A).

To visualize template switching more directly, we combined wild-type donors with recoded donors where the GFP11 insert contained several silent mutations and used Illumina sequencing to sequence the insertion en masse (Figure 7B). Using recoded donors with silent mutations every 12 bases in the GFP11 insert, we identified evidence of template switching in 1.4% of edits (“chimeric edits”, see Materials and Methods). Interestingly, the same experiment performed with donors that contained silent mutations every 6 or every 3 nucleotides resulted in only 0.5% and 0% chimeric edits, respectively (Figure 7B, Figure S5 and Table S6). The chimeric edits could not have resulted from sequential rounds of Cas9 cleavage and repair, since the edit destroyed the crRNA pairing sequence. The chimeric edits also could not have arisen during PCR amplification, since we observed no chimeric edits in a control experiment mixing two different cell populations (Figure S5). We conclude that template switching occurs between donors *in viv*o and is sensitive to the degree of homeology between donors.

### Accuracy of repair is asymmetric

To investigate the accuracy of repair with linear donors, we isolated GFP+ and GFP-cells by fluorescence-activated cell sorting from a single editing experiment targeting the *Lamin A/C* locus with a GFP-containing PCR fragment (Figure S6). Each cell was grown out as a clone and the *Lamin A/C* locus was amplified using two primers flanking the insertion site. As expected, all 48 GFP+ clones contained at least one *Lamin A/C* allele with a full-size insert (4 were homozygous with two edited alleles). We sequenced the GFP insert in 23 of the 48 GFP+ clones and identified 20 precise insertions and 3 imprecise insertions containing small in-frame indels at the left or right junction (Figures S6/S7). We also sequenced the wild-type-sized allele in 11 of the 44 heterozygous GFP+ clones, and identified 2 with wild-type sequence, 6 with indels at the DSB, and 3 with small inserts (<100 bp) corresponding to either the N-terminus or C-terminus of GFP (Figure S7). We also screened 37 GFP-clones by PCR and, surprisingly, identified 10 that contained inserts at the *Lamin A/C* locus. We sequenced 7 of the 10 inserts and identified 3 with a full-size GFP insert with out-of-frame indels at one junction and 4 with smaller GFP inserts (Figure S7).

In total, we sequenced 13 imprecise GFP edits and found only one internal deletion and one insertion in the wrong orientation (Figure S7). All other imprecise edits were full-size or truncated GFP fragments inserted in the correct orientation. All had one precise junction on the non-truncated terminus of GFP. The other junction was imprecise and contained indels (Figure S7). These observations are consistent with an asymmetric repair process that uses different mechanisms to initiate and resolve repair (see Discussion).

### mCherry-tagging of a mouse locus using a PCR donor with short homology arms

To test whether linear DNAs with short homology arms could also be used in a mammalian animal model, we designed a PCR fragment to insert mCherry in the mouse adenylyl cyclase 3 (*Adcy3*) locus. The PCR fragment contained a 739 bp insert flanked by two 36 bp homology arms designed to insert mCherry in frame near the C-terminus of *Adcy3*. The PCR fragment and *in vitro* assembled Cas9 complexes were co-injected into mouse zygotes, and the resulting pups were genotyped by PCR and Sanger sequencing (Figure S8). We identified 27/87 pups with a correct size insertion at the *Adcy3* locus (31% edit efficiency). Sequencing of 10 full-size mCherry edits revealed them all to be precise (no indels). A parallel editing experiment using an mCherry supercoiled plasmid with 500 bp HAs yielded 5 edits from 25 pups (20% edit efficiency). Similar knock-in efficiencies were also reported recently using long single-stranded donors (25).We conclude that PCR fragments with short HAs function as efficient donors in mouse embryos. PCR fragments yield edits at frequencies similar to plasmids and ssDNA donors, but with the added convenience of ease of synthesis especially for long inserts.

## Discussion

In this report, we demonstrate that PCR fragments are efficient donors for genome editing in human cells and mouse embryos. PCR fragments require only short homology arms (HAs ∼35 bp) and can be used to integrate edits up to 1kb. PCR fragments (and ssODNs) appear to participate in a replicative repair mechanism that broadly conforms to the SDSA model for gene conversion. Our findings suggest simple guidelines to streamline donor design and maximize editing efficiency (Figure S9).

### Linear DNAs repair Cas9-induced DSBs by templating repair synthesis

In principle, linear donors could repair Cas9-induced breaks by integrating directly at the DSB. For example, microhomology-mediated end-joining (MMEJ) could cause donor ends to become ligated to each side of the DSB (8). Alternatively, HAs on the donor could form holiday junctions with sequences on each side of the DSB. Cross-over resolution of the two holiday junctions could cause donor sequences to become integrated at the DSB. This type of HDR has been proposed to underlie genome editing with plasmid and viral donors (16). In these models, repair is symmetric: the same mechanism (MMEJ or recombination) is used to ligate donor sequences to each side of the break. In contrast, our observations suggest that repair with linear donors proceeds by an asymmetric, likely replicative, process. First, ssODNs with only one HA show strong polarity specificity (Figure 4), consistent with a specific requirement for pairing with 3’ ends at the DSB (Figure 1). Second, recessed HAs (HA at a distance from the DSB) are rarely used to initiate a repair event, but can be used to resolve a repair event (Figure 6). Third, internal homologies on the donor can bypass integration of distal edits (Figure 5). Fourth, most imprecise edits have asymmetric junctional signatures (Figure S7). These observations suggest that the repair process is polar like DNA synthesis and has different requirements to initiate and resolve repair. These findings are consistent with the SDSA model for gene conversion (15) (Figure 1). SDSA initiates with DNA synthesis templated by the donor to extend 3’ ends at the DSB, and resolves by annealing of the newly replicated strand(s) back to the locus. Our observations suggest that initiation of DNA synthesis is the most homology-stringent step, requiring a ∼35 base HA on the donor complementary to sequences directly adjacent to one side of the DSB. The observations that HAs longer than 35 bases do not perform significantly better, and that distal HAs perform more poorly, suggests that resection exposes only short regions of ssDNA on either side of the DSB. In contrast to the initiation step, the resolution step has more relaxed homology requirements. Recessed arms can be used for that step, and in fact repair can proceed with no HA on the “annealing side” (Figure 4). In that case, NHEJ (or MHEJ) is used to fuse the newly replicated strand to the other side of the DSB. One possibility is that NHEJ or MHEJ competes with annealing during resolution, especially in the case of long edits where synthesis has a higher chance of stalling before reaching the distal HA or before synthesis of a complementary strand primed from the other side of the DSB (Figure 1). Consistent with this view, we recovered several partial GFP insertions that were integrated in the correct orientation but contained one imprecise junction on the truncated side of GFP.

Partial edits due to premature withdrawal of the newly replicated strand from the donor should be less frequent with shorter inserts. Consistent with this prediction, we found that editing efficiency is inversely proportional to insert size. At the *Lamin A/C* locus, we obtained 45% edits for a 57 bp insert, 26% edits for 714 bp insert (GFP) and 20% edits for a 993 bp insert. The size of the insert, and not the overall size of the donor, correlated with efficiency (Figure 3). A likely possibility is that the low processivity of repair polymerases (26) increases the chances of aberrant dissociation/annealing events on long inserts.

We also obtained evidence for dissociation and invasion events between donors. Such “template switching” causes sequences from overlapping donors to become incorporated in the same edit. We found that template switching is sensitive to the degree of homology between donors and is reduced significantly by mutations every 3 or 6 bases. Similarly, recoding of sequences between the DSB and the edit can reduce the occurrence of premature annealing events that terminate synthesis on the donor before copying of the edit. Template switching may also explain why editing efficiency is sensitive to donor molarity, since high donor molarity is predicted to lower the frequency of aberrant dissociation/annealing events during synthesis. It will be interesting to determine which repair polymerases are responsible for synthesis templated by linear donors and whether their processivity characteristics account for our observations of template switching. In this regard, it is interesting to note that we identified a higher frequency of full-length edits (and lower frequency of partial edits) in mice compared to HEK293T cells. This difference could reflect differences in the properties of the enzymes that mediate SDSA in the two systems. Alternatively, the higher precision in mice could be due to a more efficient method for delivering donors at high molarity (pronuclear injection in mouse zygotes versus nucleofection in HEK293T cells).

### SDSA as a repair mechanism for Cas9-induced DSBs: implications for genome editing

The demonstration that both ssODNs and PCR fragments engage in SDSA to repair Cas9-induced DSBs in human cells has two important implications for genome editing. First, the SDSA model makes simple predictions for optimal donor design (Figure S9). These predictions improve editing efficiencies for edits at distance from the DSB, and eliminate the effort and expense used in creating donor DNAs with unnecessarily long homology arms.Linear donors with short homology arms can be chemically synthesized as single-stranded or double-stranded DNA fragments without any cloning. In this manner, tagging of genes with GFP can be achieved readily, without resourcing to split-GFP approaches that also require expression of a complementary GFP1-10 fragment (22). Second, because SDSA is thought to be a widespread mechanism for DSB repair among eukaryotes (27), it is likely that the approaches outlined here will be applicable to other cell types and organisms. We documented previously that PCR fragments with short HAs perform well in *C. elegans* (14), and we demonstrate here the same for HEK293T cells and mouse embryos. It will be interesting to investigate whether linear donors with short HAs can also be used for genome editing in pluripotent cells and post-mitotic cells.

## Acknowledgments

We thank the JHU GRCF sequencing facility, the JHU Transgenic facility and the JHU Ross Flow Cytometry Core Facility for expert support. We also thank Andrew Holland and Tyler Moyer for discussions and help with tissue culture and Boris Zinshteyn for discussions about Illumina sequencing and data analysis.This work was supported by National Institutes of Health (NIH) [grant number R01HD37047 to G.S., R01DC004553 to R.R., F32GM117814 to A.F.]. G.S. and R.G. are investigators of the Howard Hughes Medical Institute. D.H.G. is a Damon Runyon Fellow supported by the Damon Runyon Cancer Research Foundation (DRG-2280-16).

---

**Figure S1:**
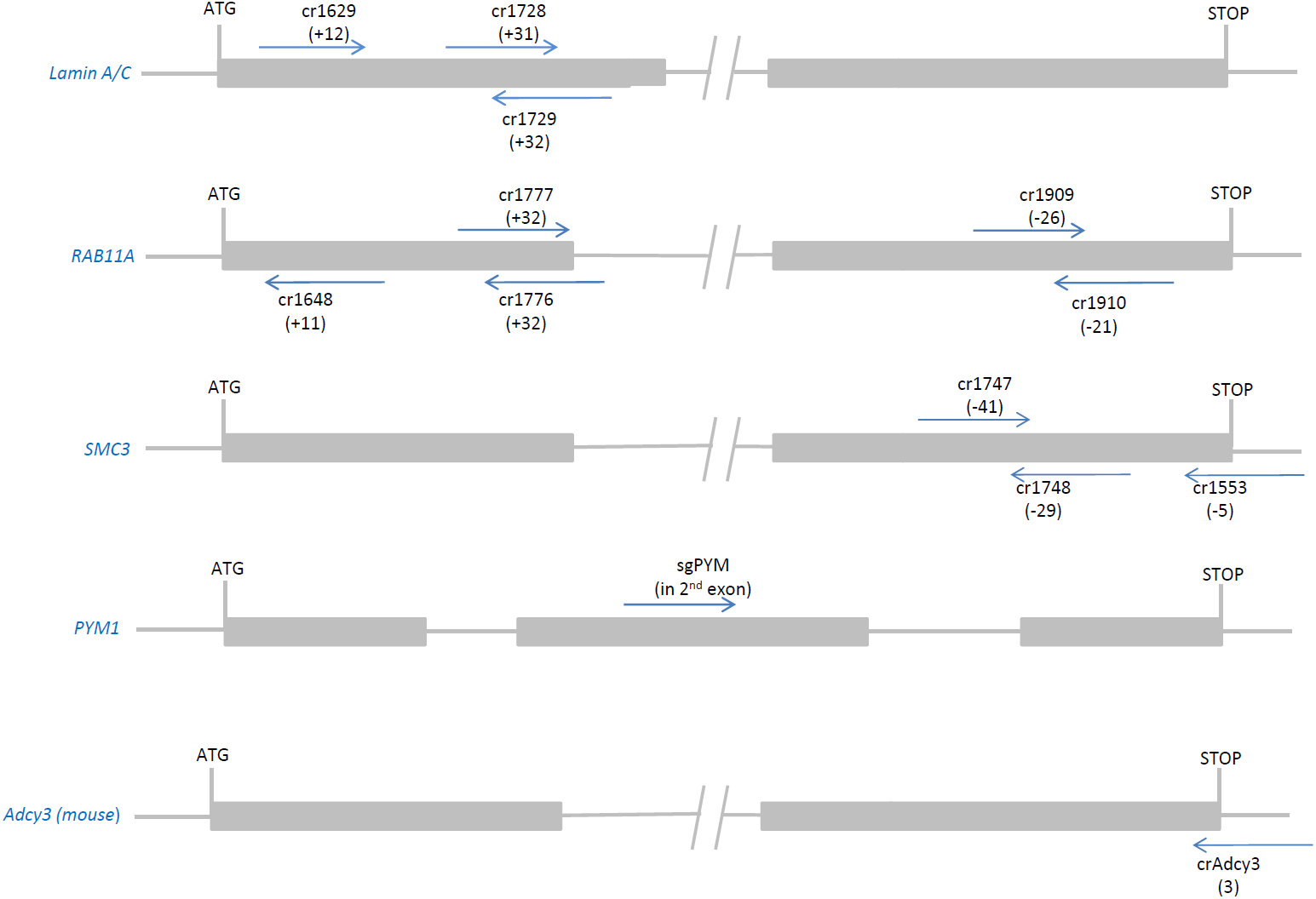
crRNAs used in this study. Schematics showing guide RNAs (arrows) used in this study mapped on *Lamin A/C, RAB11A, SMC3, PYM1* (human) and *Adcy3* (mouse) loci. Grey boxes indicate coding exons, only the first and last exons are shown for *Lamin A/C, RAB11A, SMC3*, and mouse *Adcy3*. For each guide, arrows indicate the 3’ end. Numbers indicate position of the DSB relative to the ATG or STOP codon.Chemically synthesized crRNAs were used at all loci, except for *PYM1* where we used a plasmid-encoded sgRNA. Guide RNA sequences are in Table S4.

**Figure S2:**
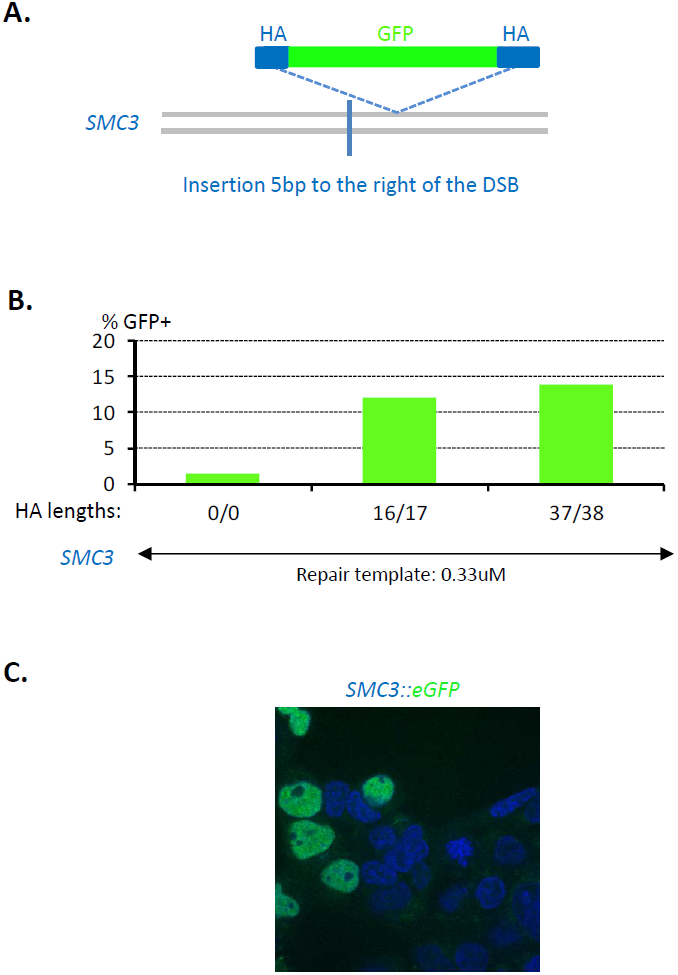
tagging with GFP of the *SMC3* locus using PCR repair template with short homology arms. **A.** Diagram showing PCR donor for GFP insertion at the *SMC3* locus. Locus - grey, GFP - green, HA (Homology Arm) - blue. GFP was inserted 5 bp to the right of the DSB. **B.** Graphs showing % of GFP+ cells obtained with PCR fragments with HAs of the indicated lengths. Insert size in all cases was 714 bp. PCR fragments were nucleofected in HEK293T cells at the concentration indicated and cells were counted by flow cytometer 3 days letter. **C.** Confocal images of cells 3 days after nucleofection. GFP: green, DNA: blue. The GFP subcellular localization is as expected for in-frame translational fusion to *SMC3*, a nuclear protein.

**Figure S3:**
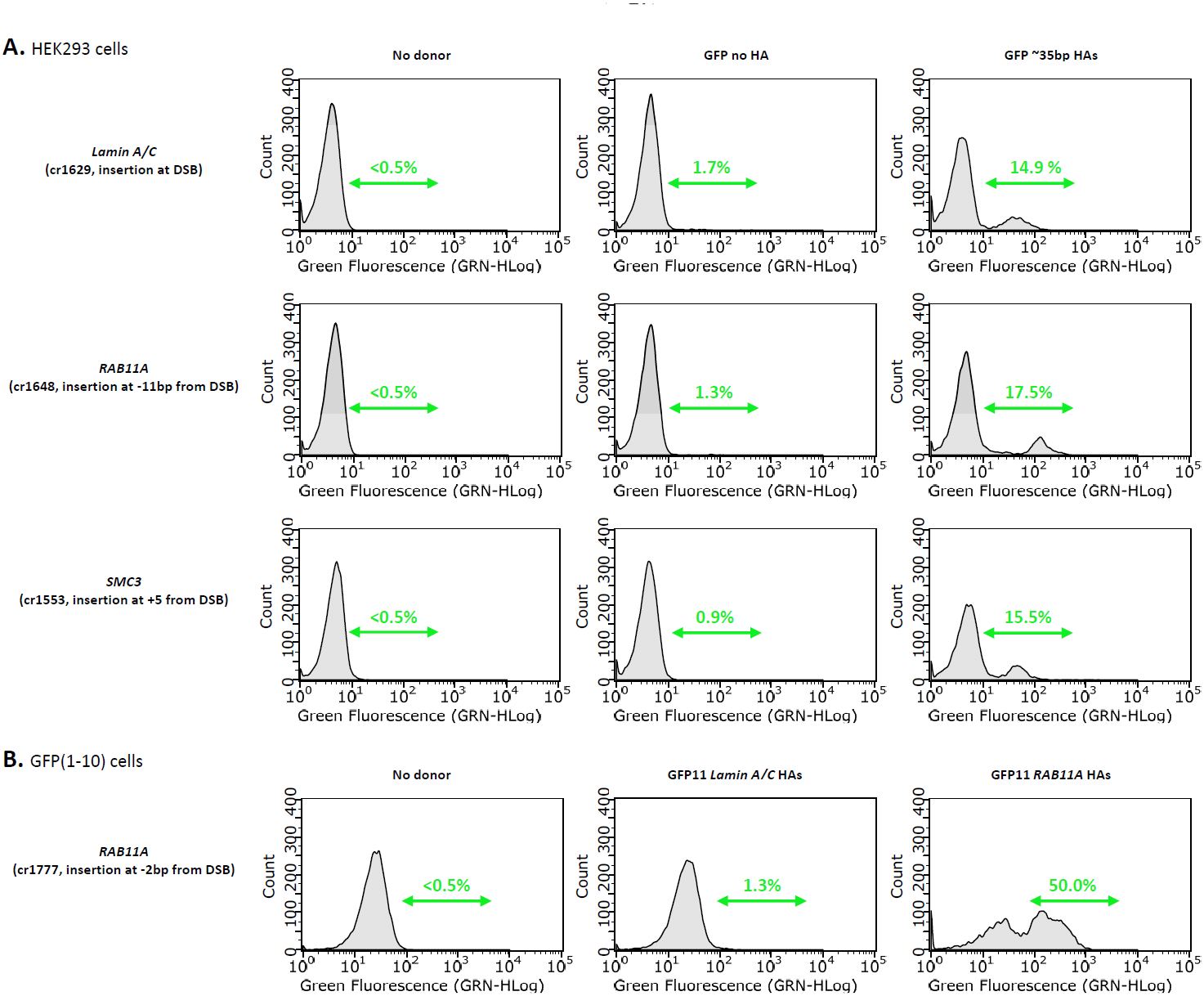
Flow cytometer plots of cells tagged with PCR repair templates. Flow cytometer plots showing the number of cells (Y axis) and their GFP intensity (X axis). **A.** *Lamin A/C, RAB11A* and *SMC3* were targeted in HEK293T cells with an eGFP containing PCR fragment with or without ∼35 bp Homology Arms (HAs). Green double arrows indicate the % of GFP+ cells. For every experiment, non-nucleofected cells were also run through the flow cytometer to determine background fluorescence (<0.5% cells). Note that donors without HAs yield GFP+ values slightly above background, consistent with a low level of integration by NHEJ or MMEJ. **B.** *RAB11A* was targeted in HEK293T (GFP1-10) cells using a GFP11-containing repair template with or without ∼35 bp Homology Arms (HAs). Green double arrows indicate the % of GFP+ cells. Non-nucleofected cells were also run through the flow cytometer to determine background fluorescence (<0.5% cells). Note that HEK293T cells that express GFP1-10 cells have a higher intrinsic fluorescence than HEK293T cells.

**Figure S4:**
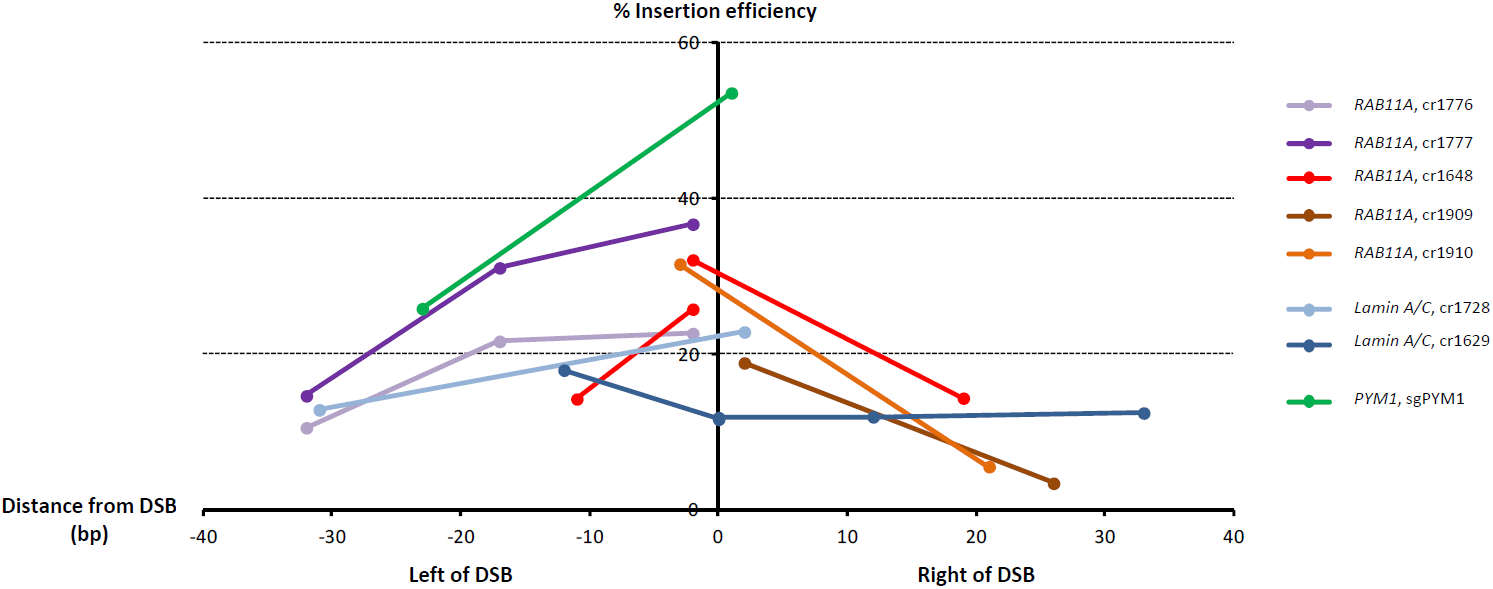
Insertion efficiency relative to distance from the DSB. Graph showing the efficiency % of editing (Y axis) *vs* distance from the DSB (X axis) (data from Figure 6). Each line links editing experiments performed with the same guide RNA. ssODNs (optimal polarity, Figure 6) were designed to insert the edit at varying distances from the DSB as indicated. For all ssODNs, the sequence between the edit and the DSB was partially recoded to minimize premature annealing and Cas9 re-cutting of the edited locus while preserving coding potential.

**Figure S5:**
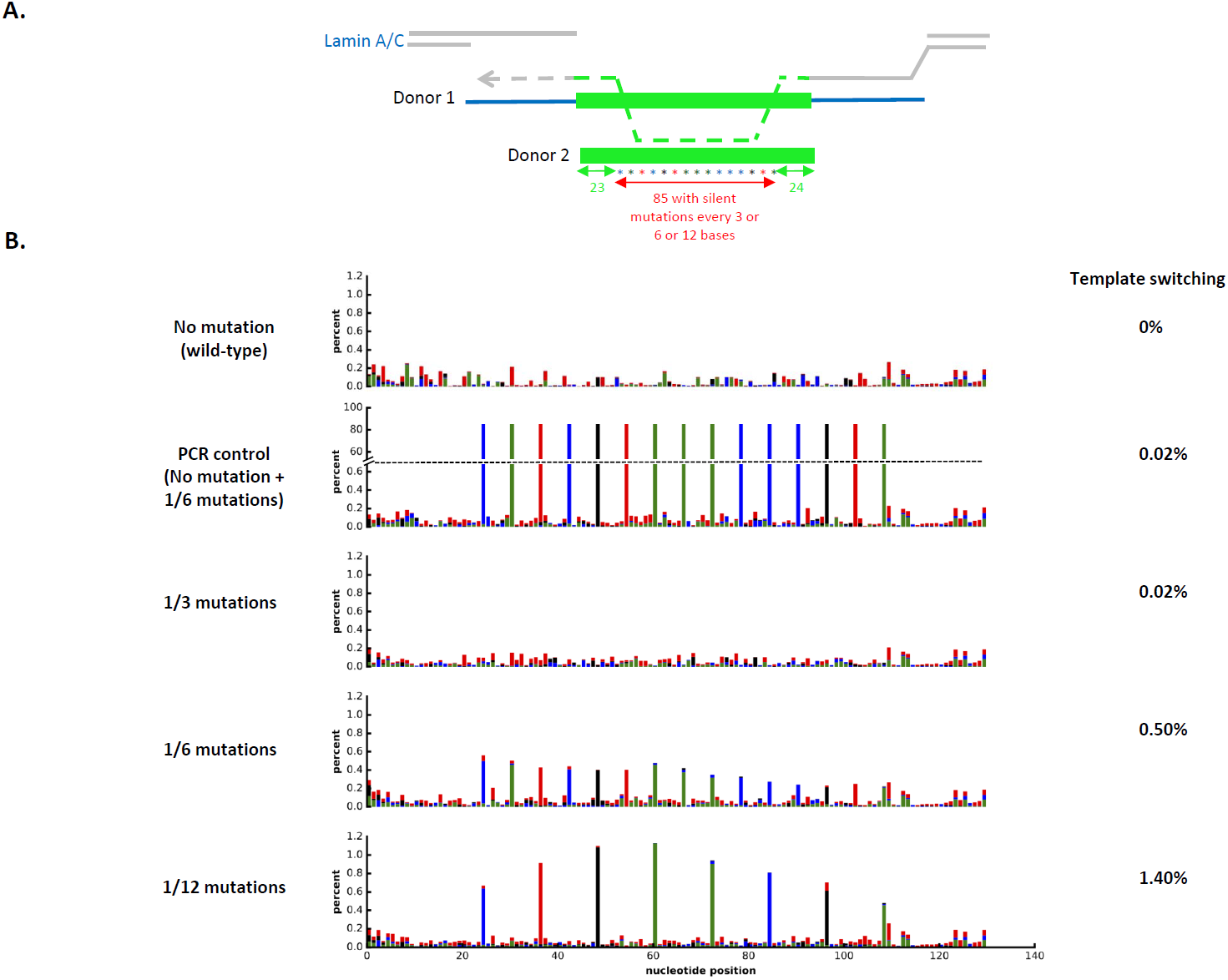
Illumina sequencing to monitor template switching. **A.** Schematic representation of the experimental design (see Figure 7). Stars in color represent silent mutations used to monitor template switching. **B.** The probability of a mutation (relative to the “No mutation” template) at each nucleotide position in the region of the repair template, after removal of incompletely mapped and low-quality reads. Bars are color-coded by identity of the incorporated nucleotide. Green: A, blue: C, black: G, red:T. PCR control: Two cell populations that received separately a wild-type ssODN or a mutant ssODN (1/6 mutations) were combined for PCR amplification. This control was used to determine basal levels of template switching that might occur during PCR amplification. These levels are 25-fold lower than observed in cells co-transfected with the same two ssODNs (0.02% versus 0.50%).

**Figure S6:**
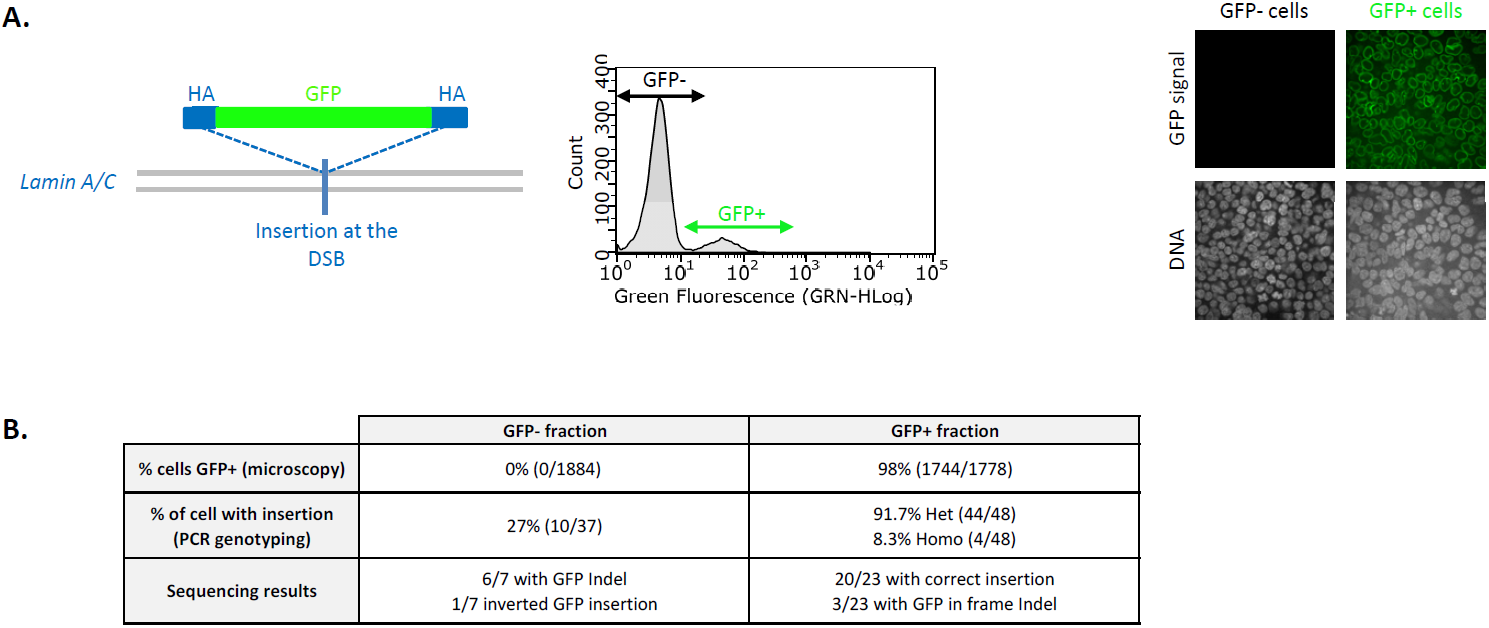
Derivation of GFP+ and GFP-clones from a single editing experiment targeting the *Lamin A/C* locus with a GFP-containing PCR fragment. **A.** Schematic showing the donor (green with blue Homology Arms - HAs) and targeted locus (grey). HEK293T cells were edited at the *Lamin A/C* locus with an eGFP PCR donor with 33/33 HAs, and FACS-sorted as GFP+ and GFP-cells. The clones were amplified and examined by confocal microscopy. All GFP+ cells exhibit the expected nuclear membrane localization expected from a GFP translation fusion with *Lamin A/C*. **B.** Statistics of genotyping results for GFP+ and GFP-single clones. See text and Figure S7 for details.

**Figure S7:**
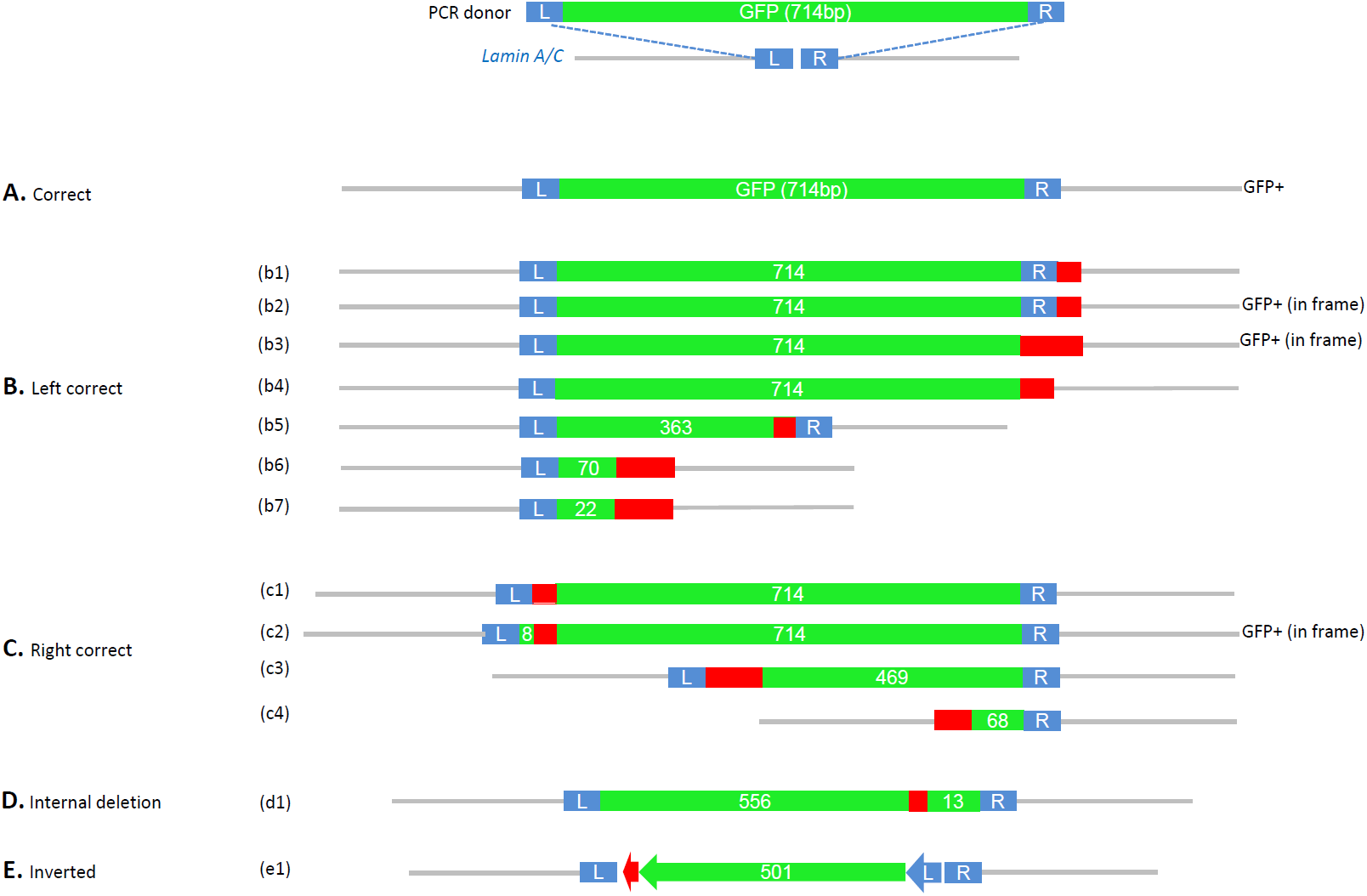
Structure of imprecise GFP knock-in edits. Schematics showing the GFP inserts obtained in the experiment described in Figure S6. *Lamin A/C* locus (grey line), Full-length left HA (L, 33 bp) and right HA (R, 33 bp) (blue), GFP (green, with length of GFP sequence indicated), Indel (red). GFP+ indicates cells with *Lamin A/C* GFP signal. **A.** Precise edit for reference **B.** Edits with imprecise right junctions (b1) Contain an 11 bp duplication of the *Lamin A/C* locus sequence just downstream the right HA. (b2) Contain a 6 bp deletion of the *Lamin A/C* locus sequence just downstream the right HA. (b3) Contain a deletion of the last 19 bp of the right HA and of the 8 bp just downstream the right HA sequence. (b4) Contain an 11 bp deletion inside the right HA. (b5) Contain only the 363 first bp of GFP sequence. (b6) Contain only the 70 first bp of GFP sequence followed by a 4 bp insertion and a full deletion of the right HA together with a 4 bp deletion of the *Lamin A/C* locus sequence just downstream the right HA sequence. Sequencing from wild-type size allele from Het GFP+ cell. (b7) Contain only the 22 first bp of GFP sequence followed by a 5 bp insertion and a deletion of the first 13 bp of the right HA. Sequencing from wild-type size allele from Het GFP+ cell. **C.** Edits with imprecise left junctions. (c1) Contain a 23 bp duplication of the left HA just upstream the GFP sequence. (c2) Contain on the left side the 8 first bp of GFP, followed by the 25 bp of the left HA sequence upstream of GFP, and followed by full-length GFP sequence. (c3) Contain a 52 bp insertion followed by the last 469 bp of GFP sequence. (c4) Contain a deletion of the last 7 bp of the left HA followed by the last 68 bp of GFP sequence. Sequencing from wild-type size allele from Het GFP+ cell. **D.** Edit with internal deletion (d1) Contain the 556 first bp of GFP sequence followed by a 12 bp insertion and the last 13 bp of GFP sequence. **E.** Edit with inverted insertion (e1) Contain the left HA and first 501 bp of GFP sequence inverted.

**Figure S8:**
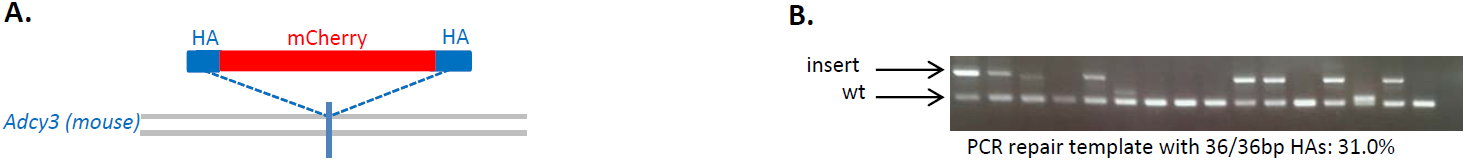
mouse *Adcy3* locus tagging with mCherry using PCR donor with short homology arms. **A.** Schematic representation of the mouse *Adcy3* locus repair strategy: mCherry (red), Homology Arms (HA, blue), locus (grey lines), DSB (blue line). B.Example of genotyping PCRs using primers flanking the DSB (outside the HAs) and run on agarose gel. The upper bands (‘insert’ arrow) correspond to the mCherry insertion. Details can be found in Materials and Methods and Table S1.

**Figure S9:**
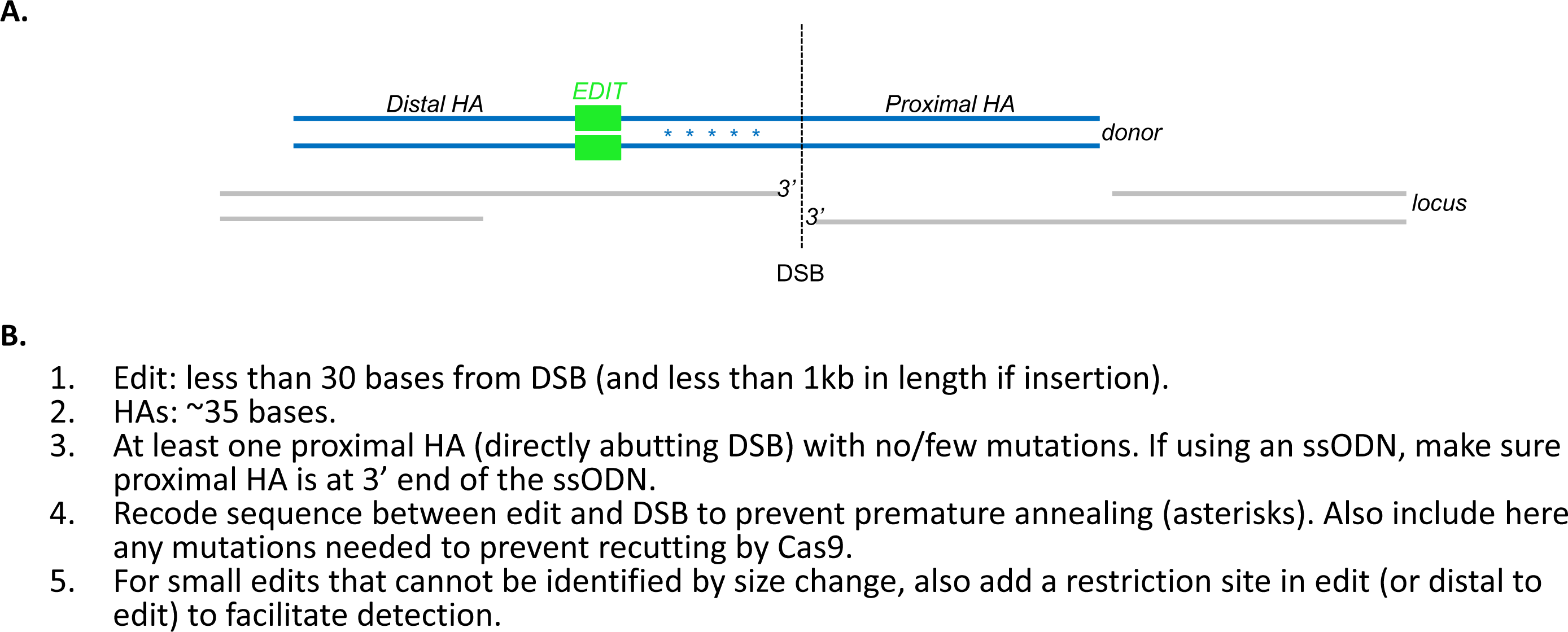
Guidelines for donor design. **A.**Schematic showing typical editing experiment to introduce an edit (green box) at a distance from the DSB (stippled line). B.Recommendations based on results presented in this study. We refer readers to (5, 21) for additional recommendations for ssODNs designed to insert edits at the DSB.

**Table S1:**
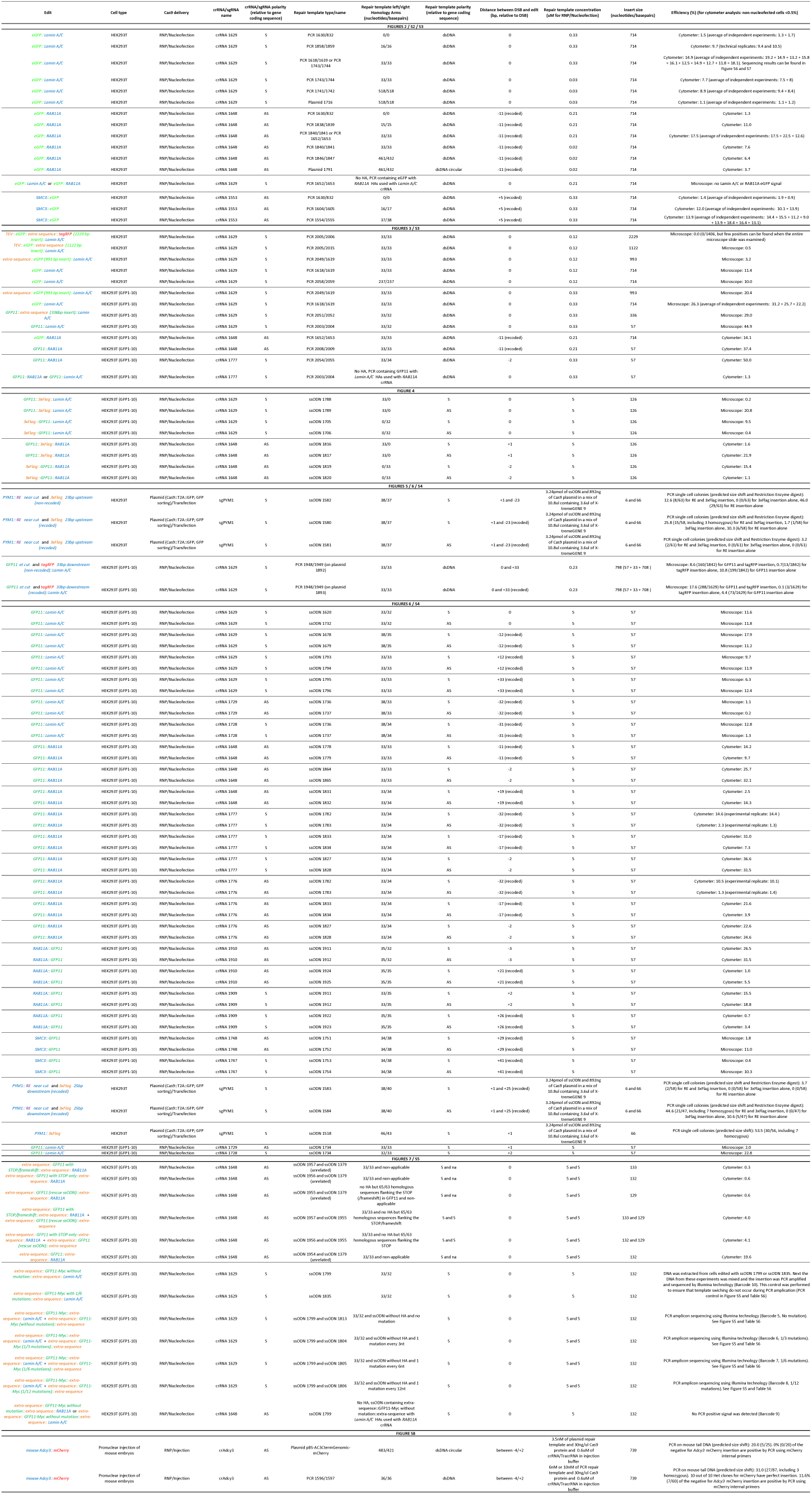
Detailed experimental conditions and results.

**Table S2:**
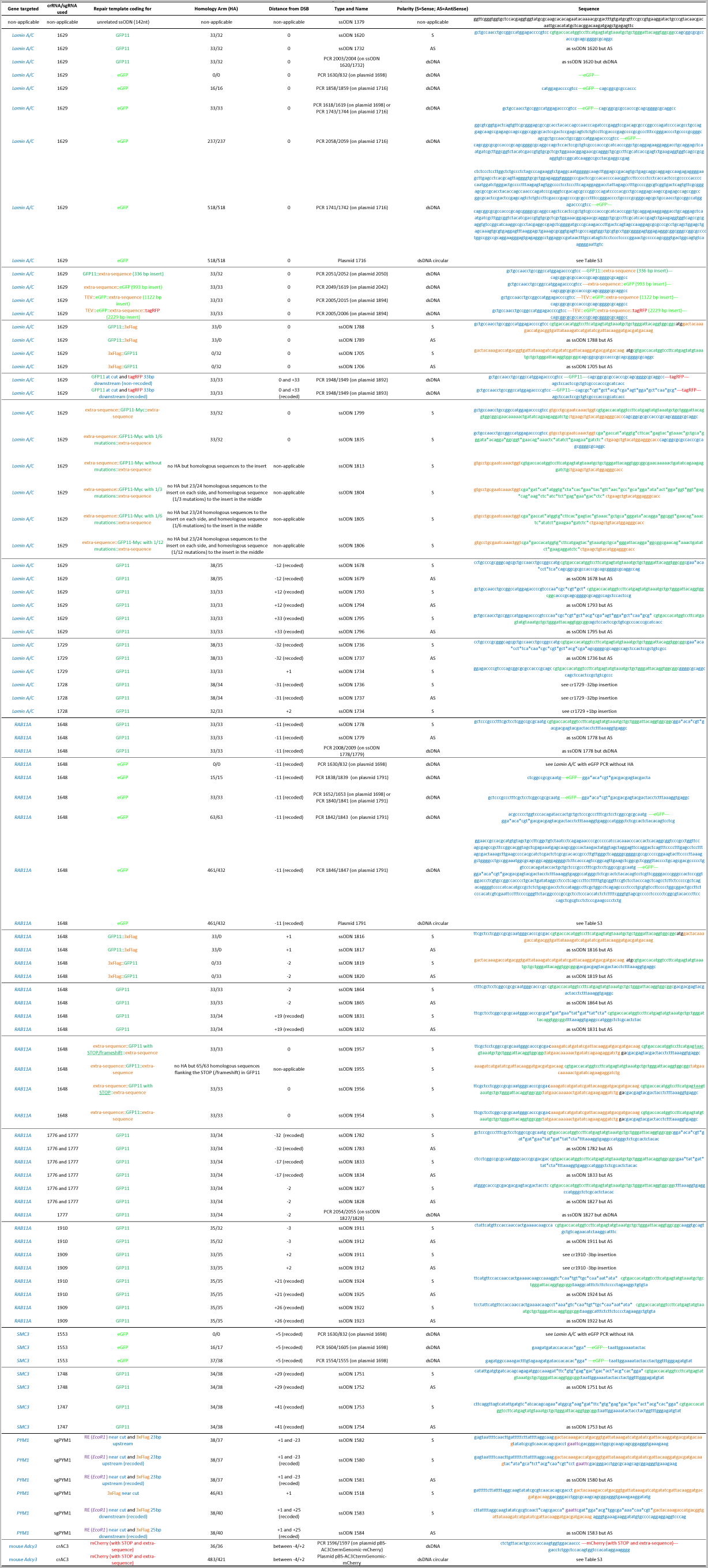
Repair templates used in this study.

**Table S3:**
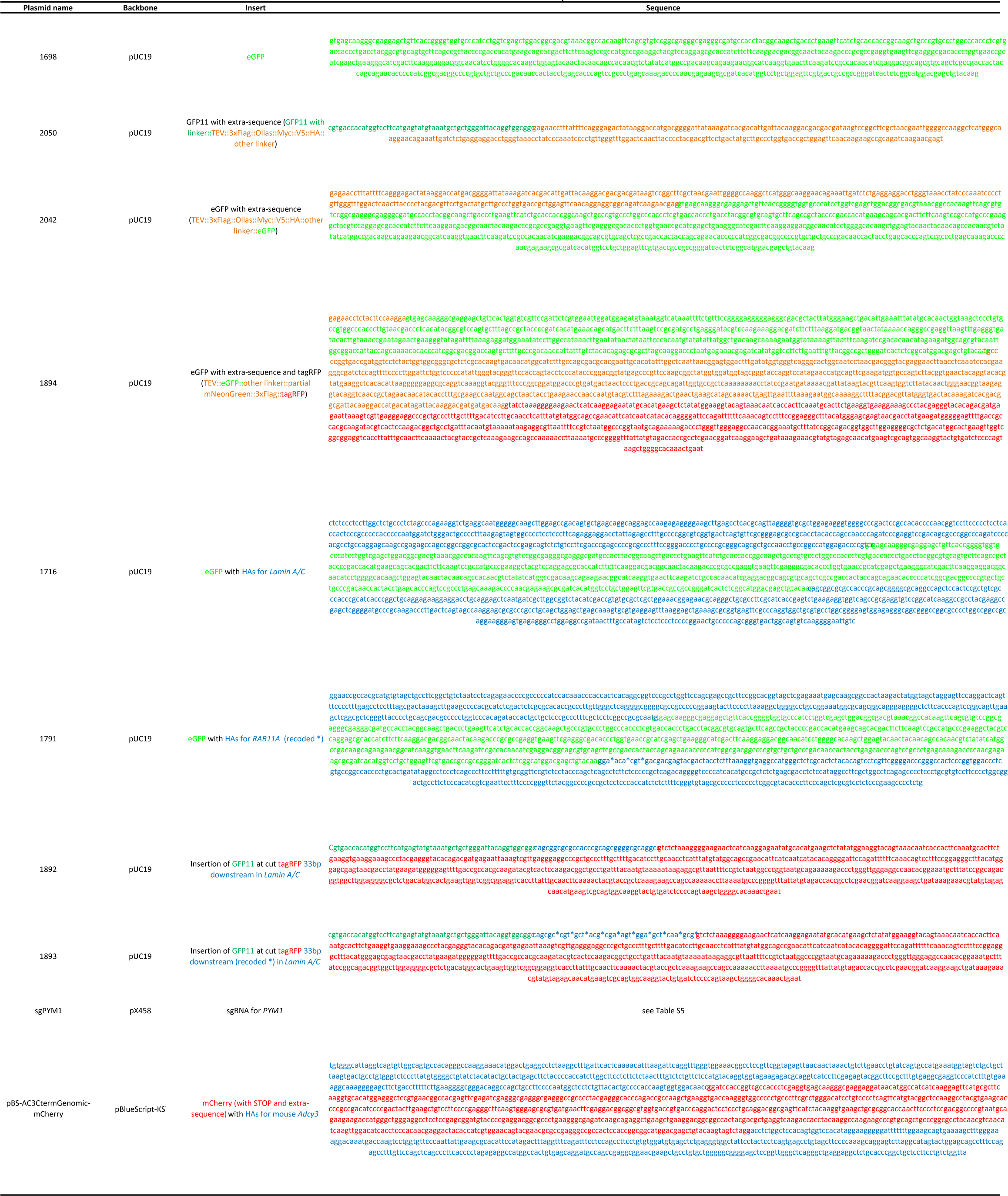
Plasmids used in this study.

**Table S4:**
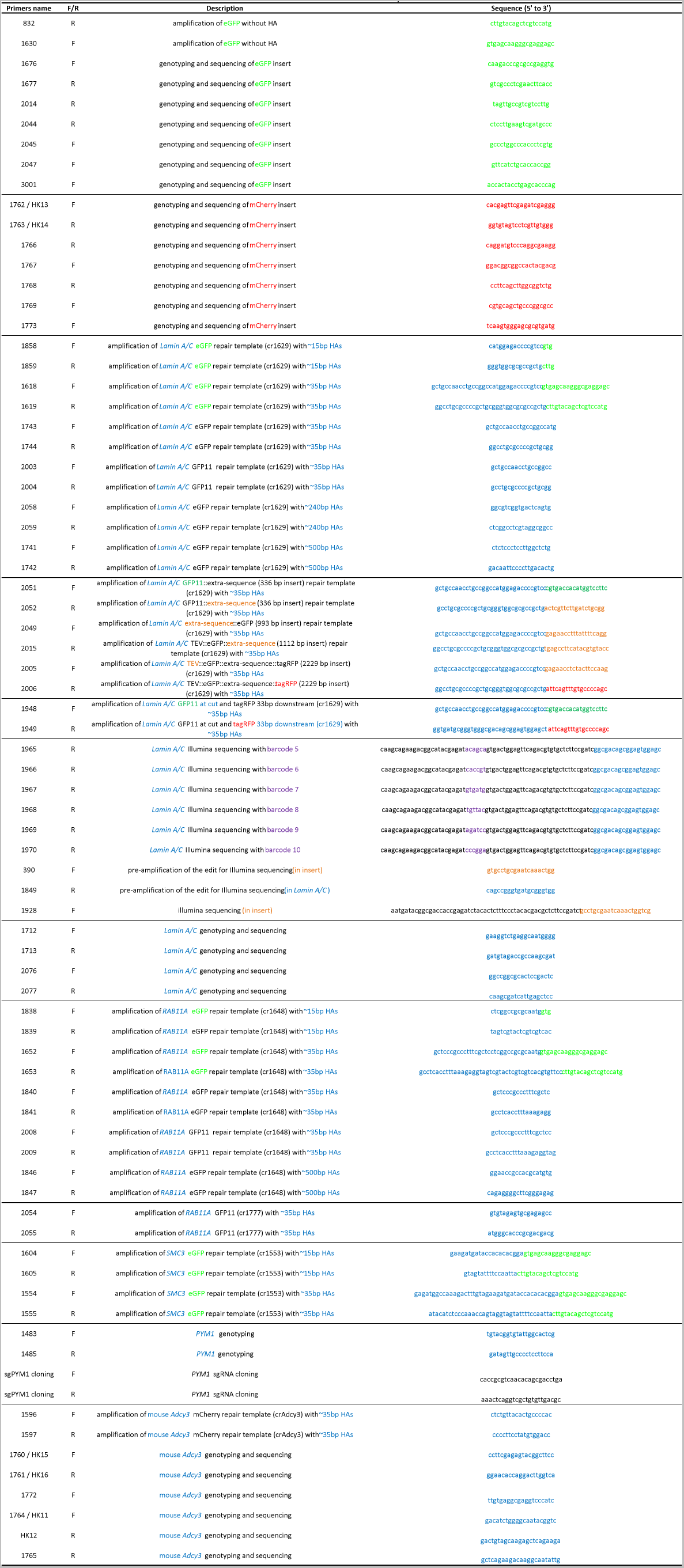
Primers used in this study.

**Table S5:**
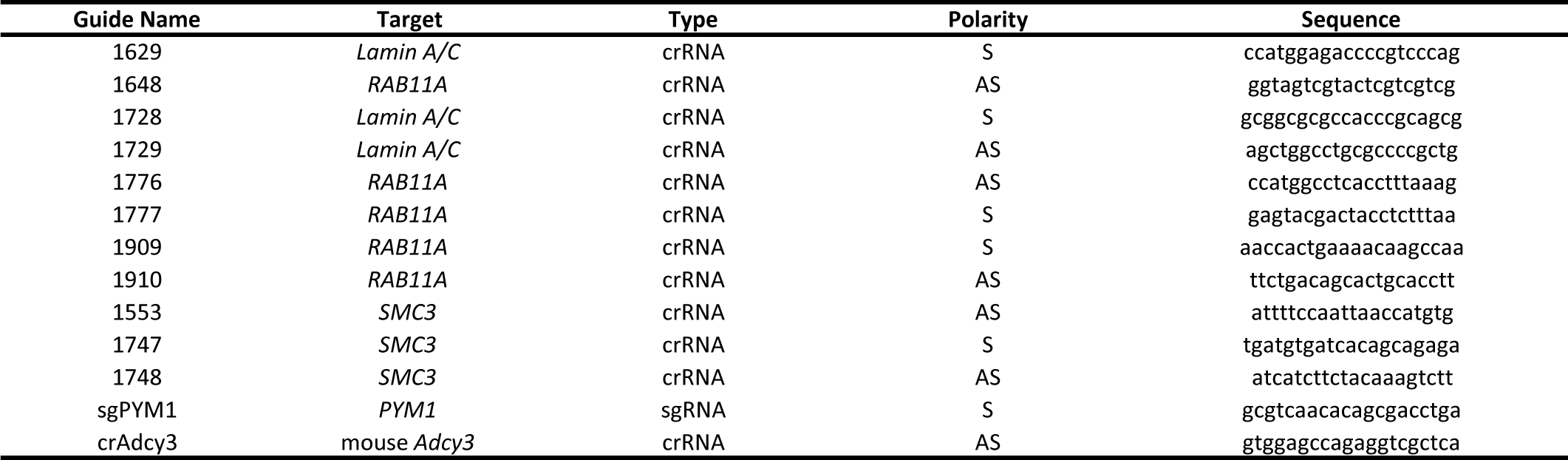
crRNA/sgRNA used in this study.

**Table S6:** Classification of reads from Illumina sequencing.

| Sample | Total sequencing reads | Do not fully map to template | Below quality threshold | Unexpected mutation at diagnostic position | Used in downstream analysis | Reads with switching detected (% of the previous column) |
| --- | --- | --- | --- | --- | --- | --- |
| No mutation | 3,369,768 | 21.60% | 42.30% | 0.20% | 36.00% | 0.00% |
| PCR control | 3,241,689 | 20.10% | 56.40% | 0.20% | 23.50% | 0.02% |
| 1/3 | 3,411,796 | 20.80% | 42.30% | 0.40% | 36.60% | 0.02% |
| 1/6 | 5,680,820 | 21.20% | 42.40% | 0.20% | 36.20% | 0.50% |
| 1/12 | 6,414,459 | 21.00% | 42.40% | 0.10% | 36.50% | 1.40% |

## References

1. Jasin M & Haber JE (2016) The democratization of gene editing: Insights from site-specific cleavage and double-strand break repair. DNA Repair (Amst) 44:6–16.

2. Doudna JA & Charpentier E (2014) Genome editing. The new frontier of genome engineering with CRISPR-Cas9. Science 346(6213):1258096.

3. Danner E, et al. (2017) Control of gene editing by manipulation of DNA repair mechanisms. Mamm Genome.

4. Liang X, Potter J, Kumar S, Ravinder N, & Chesnut JD (2017) Enhanced CRISPR/Cas9-mediated precise genome editing by improved design and delivery of gRNA, Cas9 nuclease, and donor DNA. J Biotechnol 241:136–146.

5. Richardson CD, Ray GJ, DeWitt MA, Curie GL, & Corn JE (2016) Enhancing homology-directed genome editing by catalytically active and inactive CRISPR-Cas9 using asymmetric donor DNA. Nat Biotechnol 34(3):339–344.

6. Paquet D, et al. (2016) Efficient introduction of specific homozygous and heterozygous mutations using CRISPR/Cas9. Nature 533(7601):125–129.

7. He X, et al. (2016) Knock-in of large reporter genes in human cells via CRISPR/Cas9-induced homology-dependent and independent DNA repair. Nucleic Acids Res 44(9):e85.

8. Yao X, et al. (2017) CRISPR/Cas9 - Mediated Precise Targeted Integration In Vivo Using a Double Cut Donor with Short Homology Arms. EBioMedicine 20:19-26.

9. Yamamoto Y, Bliss J, & Gerbi SA (2015) Whole Organism Genome Editing: Targeted Large DNA Insertion via ObLiGaRe Nonhomologous End-Joining in Vivo Capture. G3 (Bethesda) 5(9):1843–1847.

10. Suzuki K, et al. (2016) In vivo genome editing via CRISPR/Cas9 mediated homology-independent targeted integration. Nature 540(7631):144–149.

11. Nakade S, et al. (2014) Microhomology-mediated end-joining-dependent integration of donor DNA in cells and animals using TALENs and CRISPR/Cas9. Nat Commun 5:5560.

12. Yao X, et al. (2017) Homology-mediated end joining-based targeted integration using CRISPR/Cas9. Cell Res 27(6):801–814.

13. Zhang JP, et al. (2017) Efficient precise knockin with a double cut HDR donor after CRISPR/Cas9-mediated double-stranded DNA cleavage. Genome Biol 18(1):35.

14. Paix A, Schmidt H, & Seydoux G (2016) Cas9-assisted recombineering in C. elegans: genome editing using in vivo assembly of linear DNAs. Nucleic Acids Res.

15. Paques F & Haber JE (1999) Multiple pathways of recombination induced by double-strand breaks in Saccharomyces cerevisiae. Microbiol Mol Biol Rev 63(2):349–404.

16. Kan Y, Ruis B, Takasugi T, & Hendrickson EA (2017) Mechanisms of precise genome editing using oligonucleotide donors. Genome Res.

17. Bothmer A, et al. (2017) Characterization of the interplay between DNA repair and CRISPR/Cas9-induced DNA lesions at an endogenous locus. Nat Commun 8:13905.

18. Paix A, Folkmann A, & Seydoux G (2017) Precision genome editing using CRISPR-Cas9 and linear repair templates in C. elegans. Methods 121-122:86-93.

19. Paix A, Folkmann A, Rasoloson D, & Seydoux G (2015) High Efficiency, Homology-Directed Genome Editing in Caenorhabditis elegans Using CRISPR-Cas9 Ribonucleoprotein Complexes. Genetics 201(1):47–54.

20. Moyer TC & Holland AJ (2015) Generation of a conditional analog-sensitive kinase in human cells using CRISPR/Cas9-mediated genome engineering. Methods Cell Biol 129:19–36.

21. DeWitt MA, Corn JE, & Carroll D (2017) Genome editing via delivery of Cas9 ribonucleoprotein. Methods 121-122:9–15.

22. Leonetti MD, Sekine S, Kamiyama D, Weissman JS, & Huang B (2016) A scalable strategy for high-throughput GFP tagging of endogenous human proteins. Proc Natl Acad Sci U S A 113(25):E3501–3508.

23. Kamiyama D, et al. (2016) Versatile protein tagging in cells with split fluorescent protein. Nat Commun 7:11046.

24. Langmead B & Salzberg SL (2012) Fast gapped-read alignment with Bowtie 2.Nat Methods 9(4):357–359.

25. Quadros RM, et al. (2017) Easi-CRISPR: a robust method for one-step generation of mice carrying conditional and insertion alleles using long ssDNA donors and CRISPR ribonucleoproteins. Genome Biol 18(1):92.

26. Parsons JL, Nicolay NH, & Sharma RA (2013) Biological and therapeutic relevance of nonreplicative DNA polymerases to cancer. Antioxid Redox Signal 18(8):851–873.

27. Iyama T & Wilson DM, 3rd (2013) DNA repair mechanisms in dividing and non-dividing cells. DNA Repair (Amst) 12(8):620–636.

